# Longitudinal analysis of body weight reveals homeostatic and adaptive traits linked to lifespan in diversity outbred mice

**DOI:** 10.1101/2024.06.13.598774

**Authors:** G.V. Prateek, Zhenghao Chen, Kevin Wright, Andrea Di Francesco, Vladimir Jojic, Gary A. Churchill, Anil Raj

**Affiliations:** Calico Life Sciences LLC, South San Francisco, United States; The Jackson Labs, Bar Harbor, United States

## Abstract

Dense temporal measurements of physiological health using simple and consistent assays are essential to characterize biological processes associated with aging and evaluate the effectiveness of interventions on these processes. We measured body weight in 960 genetically diverse mice, every 7-10 days over the full course of their lifespan. We used a state space model to characterize the trajectories of body weight throughout life and derived 10 novel traits capturing the dynamics of body weight that are both associated with lifespan and heritable. Genetic mapping of these traits identified 12 genomic loci, none of which were previously mapped to body weight. We observed that the ability to stabilize body weight, despite fluctuations in energy intake and expenditure, is positively associated with lifespan and mapped to a genomic locus linked to energy homeostasis. Our results highlight the importance of dense longitudinal measurements of physiological traits for monitoring health and aging.

## Introduction

Aging is characterized by the progressive loss of physiological integrity (López-Otín et al. (2013); Freund (2019); Chen et al. (2022)). Homeostasis, the ability to maintain physiological integrity in response to intrinsic and extrinsic changes, is considered to be an important determinant of aging (Cannon (1929); Moldakozhayev and Gladyshev (2023)). Longitudinal phenotyping throughout the organism’s lifespan enables us to measure homeostasis by quantifying temporal relationships in one or more physiological traits, and associate these measures to healthspan and lifespan. In general, non-invasive procedures are required for longitudinal studies, as they allow for more frequent measurements with minimal stress to the organism. A variety of technologies can be used to collect longitudinal data with high temporal density, including body weight and composition, blood pressure, heart rate, sleep patterns, and frailty (Palliyaguru et al. (2021); Kuo et al. (2022)). Longitudinal data collected at high temporal density requires specialized computational tools to capture the time dynamics of physiological changes.

Body weight can be longitudinally measured at high frequency and changes in body weight have been linked to age-related changes in metabolism, disease outcomes, response to stress, and lifespan (Goodrick et al. (1990); Lissner et al. (1991); Bou Sleiman et al. (2022)). Body weight is influenced by a complex interplay of genetic and environmental factors (Wright et al. (2022)), and one such environmental factor is diet (Yang et al. (2014)). Over-consumption of calories and unhealthy diets can increase the risk of obesity and related diseases (Dixon (2010)). Conversely, a nutritious diet can help maintain healthy body weight and reduce the risk of age-related diseases (Willett et al. (2019)). Studies in model organisms have shown that certain dietary interventions, such as caloric restriction (CR) and intermittent fasting (IF), can extend lifespan and have beneficial health effects such as reduced inflammation, improved metabolic function, and increased cellular repair and regeneration (Fontana et al. (2010); Di Francesco et al. (2018); Longo and Anderson (2022)). However, it is unclear how diet, age and genetics influence the dynamics of body weight trajectories measured throughout life and whether summaries describing these dynamics are associated with lifespan.

We measured the body weight of 960 genetically diverse mice from weaning until death and characterized the dynamics of this classic complex trait throughout life. We developed a computational model for these body weight trajectories and derive novel phenotypes from the dynamics of body weight throughout life. We demonstrated the utility of this model to capture important functional changes that occur with aging such as the ability to maintain stable body weight in response to stress. Additionally, we examined how diet modulates these derived phenotypes as mice age, and explored whether these phenotypes are useful predictors of lifespan. Finally, we identified genetic determinants of these body weight-derived traits and quantified their heritability under different diets. We expect that body weight measured at high temporal density can reveal patters of change that are associated with longevity and healthy aging.

## Results

### Outline of study design

Body weights of mice can be easily and accurately measured. They can be altered by diets, diseases, aging, stressors, and environmental conditions. We enrolled 960 diversity outbred (DO) female mice derived from eight inbred founder strains (AJ, B6, 129, NOD, NZO, CAST, PWK, and WSB) (Svenson et al. (2012)), and used the longitudinally measured body weight, sampled at every 7-10 days over the full course of their lifespan (Francesco et al. (2023)). From enrollment (3 weeks) until 6 months of age, all mice were fed *ad libitum* (AL) on a diet of standard rodent chow (5KOG LabDiet). At 6 months of age, randomly assigned to groups of mice were switched to dietary treatments (192/group): *ad libitum* feeding (AL), 20% calorie restriction (20), 40% calorie restriction (40), fasting 1 day per week (1D), and fasting 2 consecutive days per week (2D) (**Figure 1A**). Individual body weight measures were taken approximately once a week (**Methods**).

**Figure 1:**
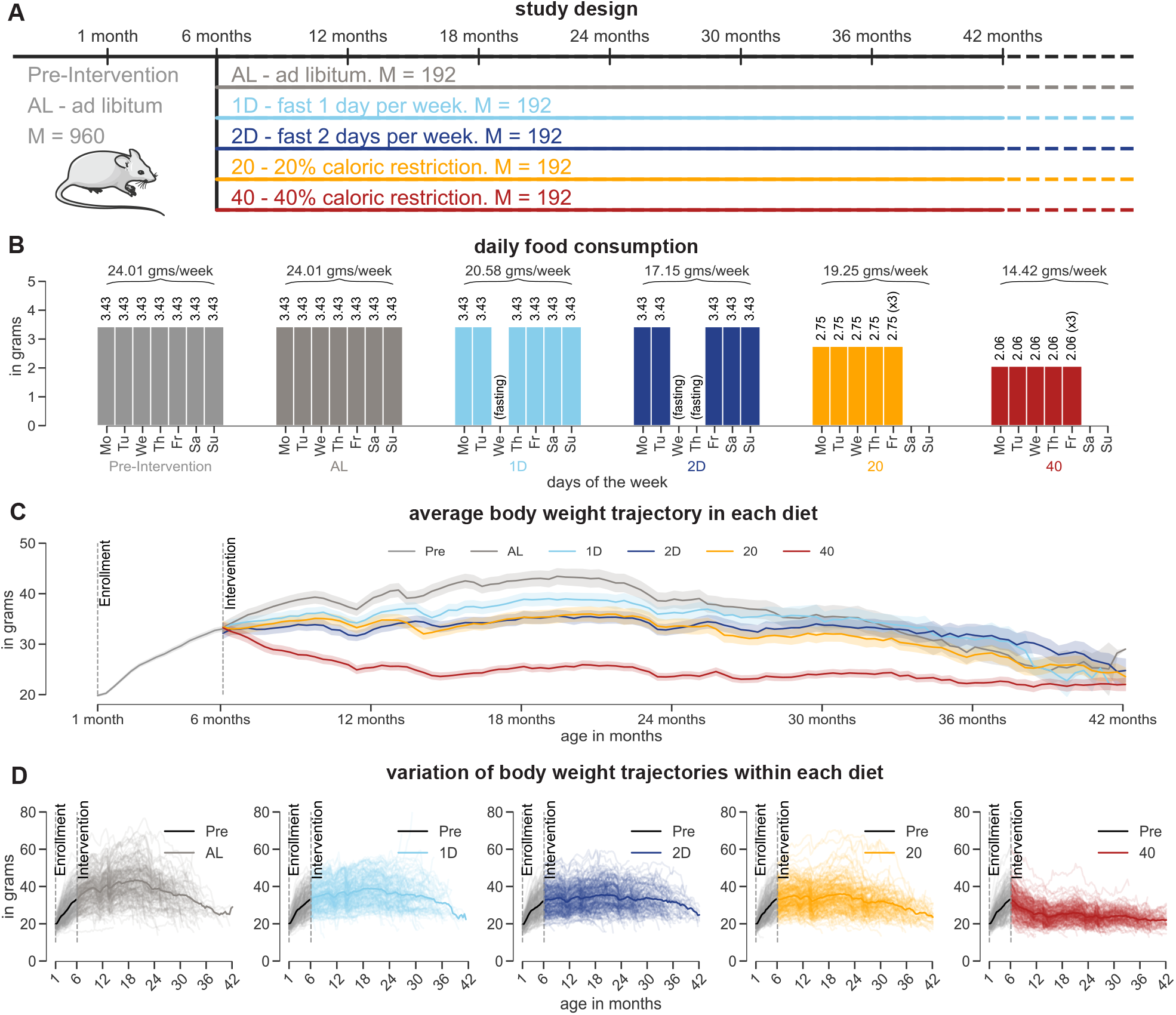
Overview of the study design. (A) At six months age, mice were randomly divided into equal proportions of five dietary groups: ad libitum (AL), one-day fasting (1D), two-day fasting (2D), 20% calorie restricted (20), and 40% calorie restricted (40). Prior to the dietary intervention, mice were on ad libitum diet. (B) Average daily food consumption (in grams) per day of the week in the pre- and post-intervention phases across diet groups. Mice belonging to the 20% and 40% groups were given three times (3x) their daily average intake every Friday. (C) Average body weight (solid lines) and standard errors (shaded region) of mice under different dietary interventions. (D) Body weight trajectories of mice in each dietary group in the pre- and post-intervention phases. The colored solid line represents the average body weight trajectory of mice in each diet.

For mice on the 1D and 2D IF diets, fasting was imposed weekly from Wednesday noon to Thursday noon and from Wednesday noon to Friday noon, respectively. These mice had unlimited food access (similar to AL mice) on their non-fasting days (**Figure 1B**). For mice on the 20% and 40% CR diets, a predetermined amount of food of 2.75 grams/mouse/day and 2.06 grams/mouse/day, respectively, were given every afternoon from Monday to Thursday, and were triple fed (x3) on Friday afternoon to last until Monday afternoon (**Figure 1B**). The predetermined amount of food was determined from a previous internal study at the Jackson Laboratory where the average amount of food consumed by female DO mice at 6 months of age was estimated to be 3.43 grams/day.

### Modeling the dynamics of body weight

Following the onset of interventions, diet was the largest source of variation in average body weight across the groups (**Figure 1C**). However, within each diet group, we observed substantial variation across mice and ages (**Figure 1D**). In order to model the temporal variation in body weight, we developed an autoregressive hidden Markov model (ARHMM) that captures the dynamics of body weight using discrete latent physiological states (**Figure 2A**).

**Figure 2:**
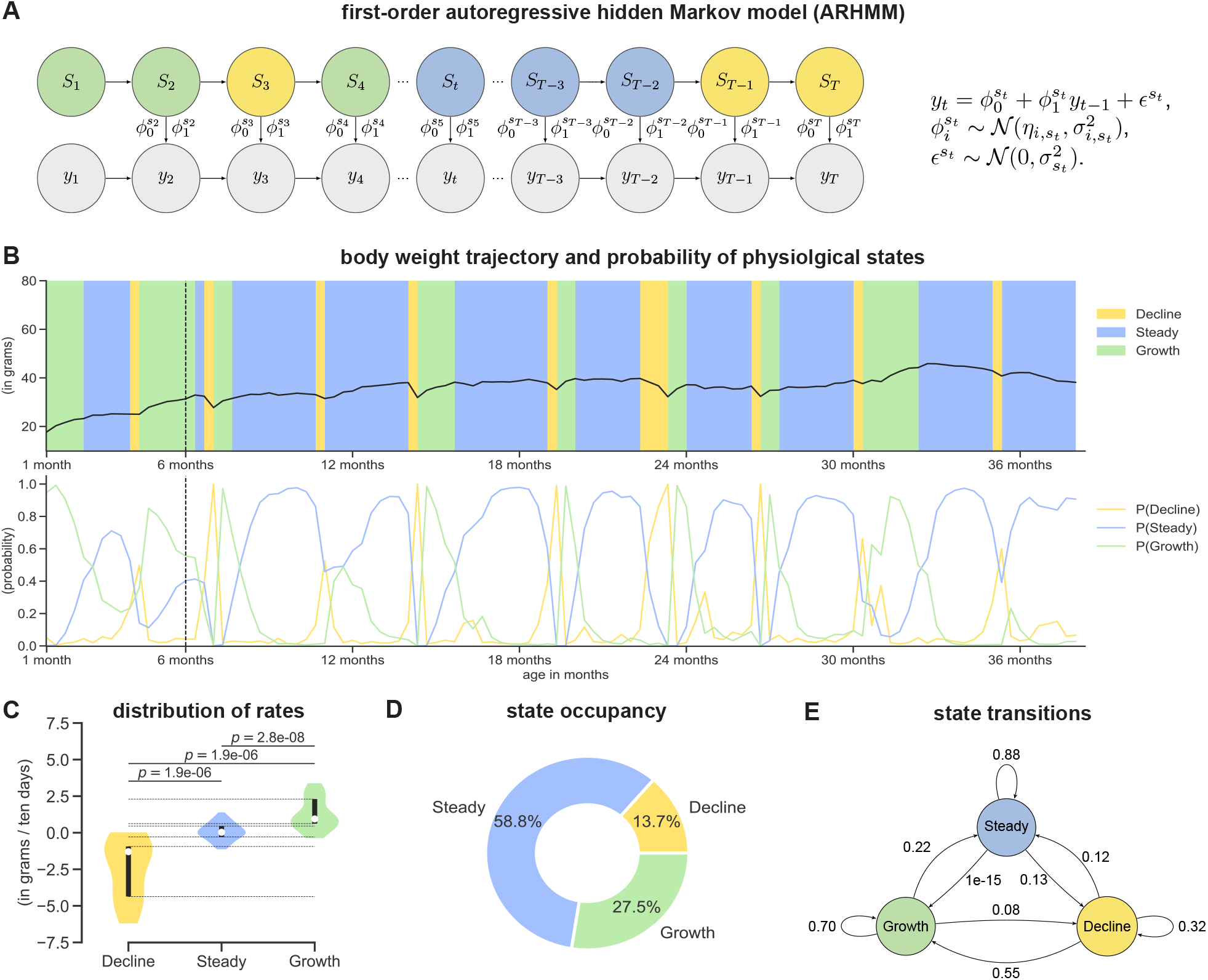
ARHMM and body weight-derived phenotypes. (A) A graphical illustration of an autoregressive hidden Markov model (ARHMM). (B) Sample body weight trajectory of a mouse on a one-day fasting diet. The start time of the intervention is indicated with a dashed vertical line. The underlying physiological states (growth, steady, and decline) are represented using background colors (top panel). The posterior probability trace of a physiological state for the corresponding body weight trajectory and its inferred dynamics (bottom panel). (C) Distribution of the rates conditioned on the physiological states. (D) The percentage of time spent in each physiological state. (E) Empirical state transition probabilities computed using the inferred states for the sample mouse.

The ARHMM combines an autoregressive model and a hidden Markov model to model a body weight trace (time-series) as a sequence of latent states. The latent states evolve according to a Markov process and capture the switching dynamics of body weight states over time. The autoregressive model captures the relationship between the current and past observations conditional on the current latent state (**Methods**). We trained the ARHMM with 70% of body weight traces from each dietary group. To determine the number of latent states of the ARHMM, we computed the deviance information criteria (Spiegelhalter et al. (2002)) on the held-out body weight traces for different numbers of latent states and starting from different random initializations (**Supplementary Figure 1A**). We obtained the smallest deviance information criteria for a model of order *K* = 3. For the given model order, the random initialization which resulted in the highest evidence-based lower bound (ELBO) on the training set was selected as the best model (**Supplementary Figures 1B-1D**). Based on the sign and magnitude of the intercept parameters for the three latent states (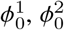, and 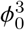), we named the states as *decline, steady*, and *growth* (**Supplementary Table 1**).

For each body weight trace, at each time point, we inferred the posterior probability that the mouse belonged to one of these three physiological states, using the best ARHMM model, and assigned the state with the highest posterior probability to that time point (**Figure 2B**). We found little overlap in the inter-quartile range of estimated rates of change of body weight within each state, indicating that these states can be well-resolved throughout the lifespan of a mouse (**Figure 2C**). Our inferences under the ARHMM model allowed us to derive several state-related traits, such as *state occupancy* and *state transitions*. For every mouse, the *state occupancy* was defined as the percentage of time the mouse spent in each state (**Figure 2D**). Similarly, the *state transition* was defined as the frequency with which the mouse switched from one state to another (**Figure 2E**). We accounted for uncertainties in state assignments when deriving these traits by using a weighted average approach, where the posterior probabilities of the state assignments were used as weights. In the next sections, we explore the effects of age, diet, and genetics on these traits and how informative they are in predicting mouse lifespan.

### Calorie restriction improves body weight homeostasis throughout life

Homeostasis is the maintenance of physiological processes in response to intrinsic and extrinsic changes experienced by an organism (Billman (2020)). Body weight homeostasis is defined as a state of unchanging body weight in response to environmental perturbations (Ravussin et al. (2014)). Modeling the dynamics of body weight allows us to generalize the definition of body weight homeostasis to include *maintenance of constant body weight dynamics*. We hypothesize that homeostasis of body weight dynamics can be influenced by a combination of various factors including diet, genetics, energy imbalance, and environmental stress.

To characterize homeostasis of body weight dynamics across diets as mice age, we divided the time-axis into six-month non-overlapping intervals starting from birth to 42 months. For each age bin, we defined *steady state homeostasis* as the fraction of time spent in the steady state (**Figure 3A**, left panel). We observed that, on average, mice spent over 60% of time post-intervention in steady state homeostasis. However, we observed large variation in steady state homeostasis across diets; on average, over a 6-month period, 40% CR mice could maintain body weight for nearly 4.5 months while AL mice could do so for just 3.5 months. When we changed the time-axis from six-month interval bins to ten percent interval bins of proportion of life lived, we observed that 40% CR mice spent nearly 75% of their time in steady state homeostasis into the last decile of life (**Figure 3B**, left panel). We could redefine steady state homeostasis in terms of the empirical probability of not transitioning out of steady state. The higher this probability, the more stable the steady state is in the mouse. We observed similar trends in the effects of age and diet on steady state homeostasis when it was redefined in terms of state stability (**Supplementary Figures 2A-2B**); left panels).

**Figure 3:**
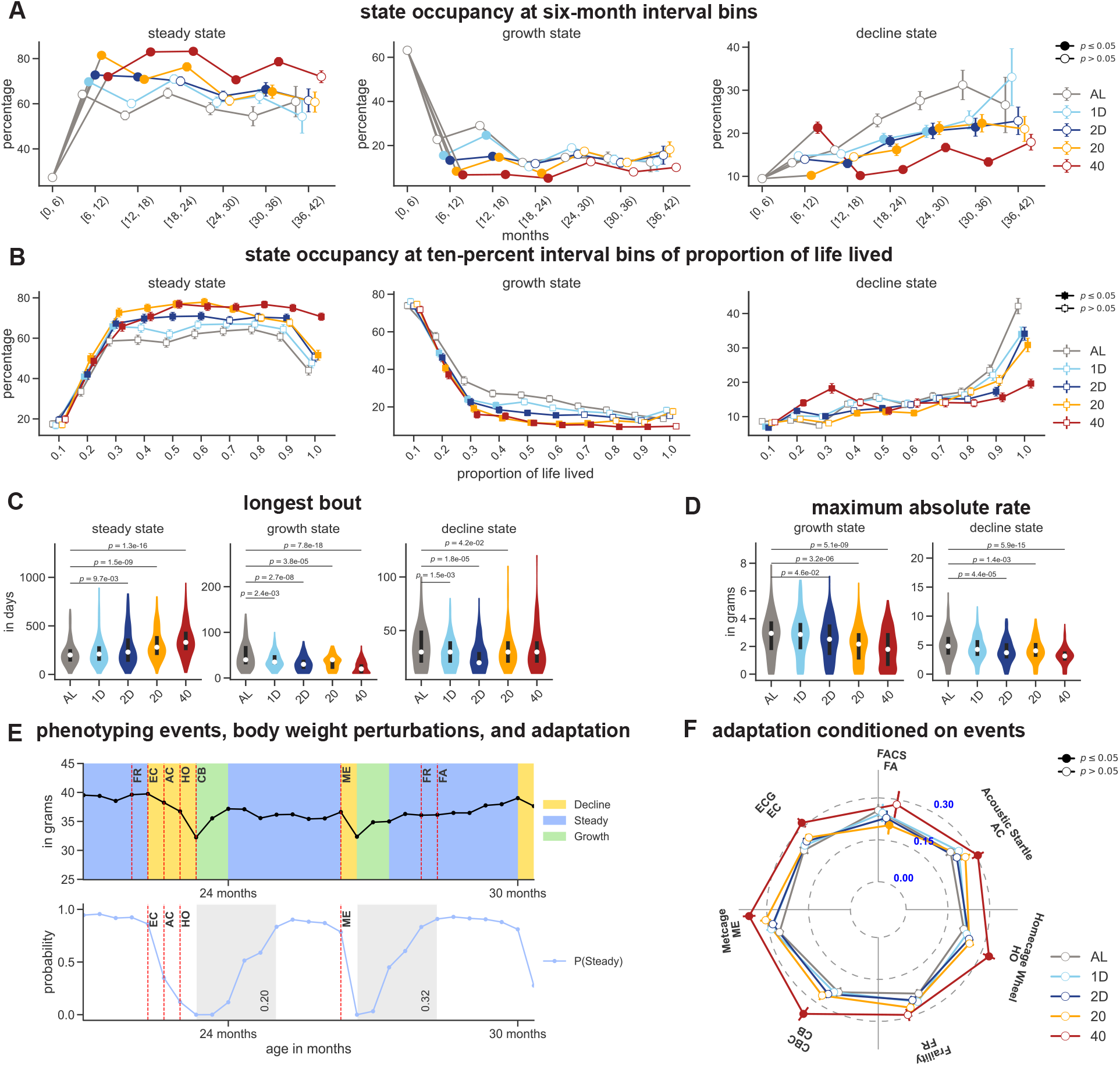
Influence of diet and age on body weight homeostasis. (A) The mean and standard errors of the state occupancy in steady, growth, and decline states, at the six-month interval bins, are represented with a circle and vertical line, respectively. The pre-intervention phase, i.e., [0, 6) months, is indicated in gray color. (B) The mean and standard errors of the state occupancy in steady, growth, and decline states, at ten percent interval bins of proportion of life lived, are represented with a square and vertical line, respectively. (C) The longest continuous bout (in days) in the growth, steady, and decline states in the post-intervention phases. (D) The maximum absolute rate (in grams) of the growth and decline states in the post-intervention phases. (E) A zoomed-in plot of the body weight trace of a sample mouse on one-day fasting diet. The approximate time of a phenotyping event is indicated using red-dashed line (top panel). The posterior probability of steady state for the corresponding body weight trace. The time period of adaptation to stress are indicated using gray boxes. (F) Radar chart of the average value and the standard-error of adaptation to stress conditioned on the phenotypic assay. In (A), (B), and (F), solid squares or circles indicate *p*-values < 0.05, where the *p*-values were obtained by performing a Mann-Whitney test between a diet group and the AL group conditioned on the same interval bin or phenotyping assay. In (C) and (D), *p*-values were obtained by performing a Mann-Whitney test between two diets, with the AL diet as the reference group.

Similar to steady state homeostasis, we defined *growth state homeostasis* as the fraction of time spent in the growth state (**Figure 3A**, center panel) and *decline state homeostasis* as the fraction of time spent in the decline state (**Figure 3A**, right panel). As expected, we observed that AL mice, on average, gained body weight for at least 4 months out of their first 6 months of life; this growth state homeostasis dropped to approximately 20% at 18 months of age. Unsurprisingly, CR and IF significantly reduced the ability of a mouse to sustain growth, with 40% CR mice spending less than 3 weeks gaining body weight over the year following dietary intervention. On the other hand, we observed that decline state homeostasis in AL mice substantially increased with age after 18 months, with mice spending nearly 30% of their time on average losing body weight until the end of their life. In contrast, calorie restriction severely reduced decline state homeostasis, with 40% CR mice spending only 10-15% of their time losing body weight after 18 months. Similar trends were also observed in the effects of age and diet on growth and decline state homeostasis when the time-axis was changed to proportion of life lived and homeostasis was defined in terms of state stability rather than state occupancy (**Supplementary Figures 2C-2F**)).

We quantified age-independent summaries of body weight trajectories post-intervention such as longest continuous bout and maximum rate of change of body weight conditioned on the latent state, and explored the effects of diet on these measures. Calorie restriction significantly increased the longest continuous bout in steady state while IF had no significant effect on this trait (**Figure 3C**, left panel). The average longest stretch of maintaining body weight lasted longer than 30% of the post-intervention lifespan for the average CR mouse compared to approximately 25% for the average AL mouse. (**Supplementary Figure 3A**, left panel). On the other hand, both CR and IF significantly shortened the longest continuous bout of gaining or losing body weight (**Figure 3C, Supplementary Figure 3A**, center and right panels). While both CR and IF affected the lifespan-normalized time of onset of the longest stretch of body weight gain, only CR affected the lifespan-normalized time of onset of the longest stretch of body weight loss (**Supplementary Figures 3B-3C**, center and right panels).

Steady state homeostasis is disrupted when mice experience a sudden change in body weight. To quantify this disruption, we calculated the maximum rate of change of body weight in the growth and decline states. AL mice achieved significantly faster post-intervention gain and loss of body weight compared to mice on other diets (**Figure 3D**). However, the effects of diet on these phenotypes were no longer significant when the rates were normalized by body weight, measured at the time when the maximum rates were recorded (**Supplementary Figure 3D**). Nonetheless, the average mouse could achieve an 8% per week increase in body weight and a 15% per week drop in body weight, suggesting that stressful life events can cause strong perturbations to body weight. However, diet did not have a substantial effect on the time at which the maximum rate of change in body weight were recorded (**Supplementary Figures 3E-3F**).

### Calorie restriction increases adaptation to stress

Mice in this study experienced experience stress due to handling during an annual one-month long phenotyping event (Francesco et al. (2023)) (**Supplementary Figure 4A**). Handling involved interactions with lab technicians and instruments, changes between group and single housing, periodic availability of running wheels, and other stressors which resulted in the disruption of steady state homeostasis (**Figure 3E**, top panel). We defined *adaptation to stress* as the rate at which a mouse returns to steady state dynamics following a deviation from steady state caused by one or more phenotyping events. We identified time windows immediately following a phenotyping event (or group of events) where a mouse is returning to steady state (shaded portions on bottom panel of **Figure 3E**), and measured the rate of adaptation to these stressor events. Specifically, we modeled the steady state posterior probability following a phenotyping event using a cumulative exponential distribution function and selected the parameter that generated the best fit to this function as the rate of return to steady state (**Methods**). The higher the adaptability to stress, the faster a mouse returns to steady state.

We measured adaptation to stress following seven of the ten phenotyping assays which substantially disrupted mice from steady state (**Supplementary Figure 4B**). Across all assays, on average, 40% CR mice showed the fastest adaptation to stress (**Figure 3F**). To quantify age-related effects on adaptation to stress, we divided the age-axis into two non-overlapping bins: (6-18) months and (18-30) months; the age-bins were picked to ensure that the number of assay events in each age-bin were equal. Across most assays, we noticed that AL mice recovered to steady state more rapidly in late life than in mid life (**Supplementary Figures 4C-4D**), while mice on other diets did not show a significant difference in stress adaptation between mid-life and late-life. This suggests that a short duration (3-4 months) of dietary restriction induced and maintained an equivalent amount of adaptability to stress as that induced by the stressors associated with a battery of phenotyping assays spread over a year.

### Steady state homeostasis is associated with reduced mortality throughout life

Body weight in mice is known to be predictive of lifespan in an age-dependent manner (Goodrick et al. (1990)); however, it is unknown whether the dynamics of body weight at different ages are also predictive of lifespan. We estimated the age-dependent effect of our derived homeostatic traits on mortality hazard (**Figure 4A**), after controlling for the effects of other lifespan determinants such as diet and body weight. Specifically, we applied a time-varying Cox proportional hazard model (Zhang et al. (2018)) to describe the survival time as a function of time-independent covariates such as diet and generation, and time-dependent covariates such as body weight (densely measured) and age-binned measures of homeostasis (sparsely quantified) (**Methods**).

**Figure 4:**
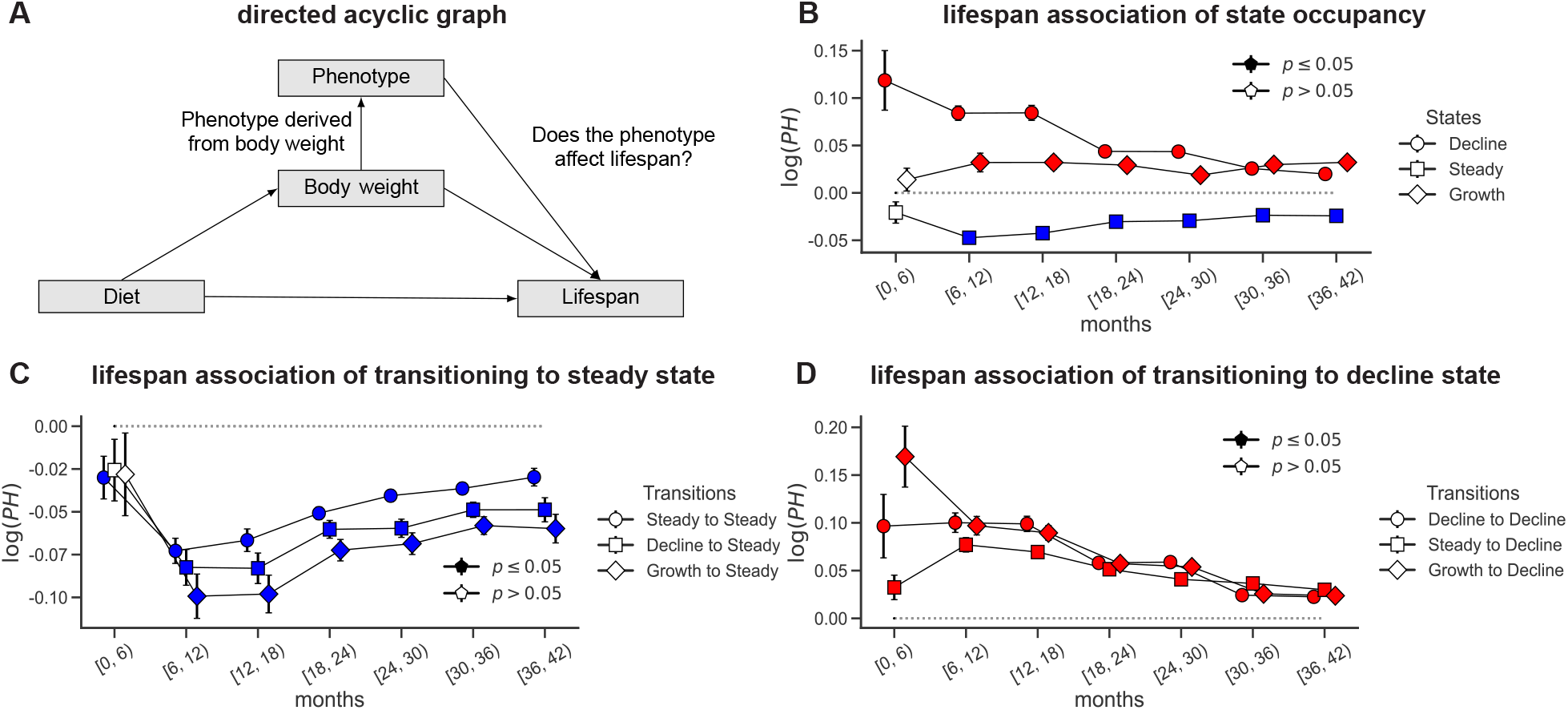
Influence of body weight homeostasis on lifespan. (A) A directed acyclic graph capturing the dependencies between covariates and lifespan. We used a time-varying Cox proportional hazard model to determine the association between body weight-derived traits and lifespan, while also accounting for confounding factors such as body weight and diet. (B) Effect size and standard error of state occupancy in steady, growth, and decline states, and its association with lifespan at every six-month interval bins. (C) Effect size and standard error of transitioning to steady state and its association with lifespan at every six-month interval bins. (D) Effect size and standard error of transitioning to decline state and its association with lifespan at every six-month interval bins. In (B), (C), and (D), lifespan associations which were statistically significant (*p*-value < 0.05) were indicated with solid colors. If the effect size of the trait at a given time interval lies above the dashed gray line (red filled markers), then an increase in the value of the trait decreases lifespan. Conversely, if it lies below the dashed gray line (blue filled markers), then an increase in the value of the trait increases lifespan.

We observed that steady state homeostasis was positively associated with lifespan, the strength of association was highest in mid-life (6-18 months) and gradually decreased with age (**Figure 4B**). Conversely, decline state homeostasis and growth state homeostasis were both negatively associated with lifespan, with a much stronger effect for decline state homeostasis. Although mice on average spent only 2 weeks in the decline state in early life (0-6 months) (**Figure 3C**), the association with lifespan was strongest for this age bin, highlighting that loss of body weight during early life in response to stressful stimuli has severe detrimental effects on lifespan.

Because increased steady state homeostasis was associated with increased lifespan, we sought to quantify the effect of transitioning to steady state (from other states) on lifespan. We observed that the rate of transitioning to steady state was positively associated with lifespan as well, although the effect sizes were dependent on the physiological state from which the mouse transitioned (**Figure 4C**). Similar to our earlier observation, high rates of transitioning to decline and growth states were associated with a decreased lifespan (**Figure 4D, Supplementary Figure 5A**). Additionally, transitions from growth to decline during early life showed the strongest negative effect on lifespan (**Figure 4D**), highlighting that perturbations to steady growth in early life that result in a loss of body weight are severely detrimental to mortality. In contrast to our homeostatic traits, adaptation to stress due to the phenotyping events showed no significant association with lifespan (**Supplementary Figure 5B**).

### Genetic determinants of body weight dynamics are distinct from those of body weight

We carried genetic mapping of our derived body weight traits on the DO mice using the GxEMM model (Wright et al. (2022)). We tested for the additive effects of genotyped variants on our derived homeostatic traits (**Supplementary Table 2**). We computed these traits using the entire post-intervention phase and focused on 18 traits that were nominally associated with lifespan (*p*-value < 0.05) (**Supplementary Figures 5C-5D**). Grouping the traits by their direction of effect on lifespan, we identified 10 traits with at least one significant variant following group-specific false discovery rate (FDR) correction, and a total of 12 loci significantly associated with these traits (**Figure 5A**; see (**Methods**) for details). None of these loci overlapped with genetic loci previously mapped to body weight of mice from the same experiment (Wright et al. (2022)). We tested for additive genetic effects on 7 lifespan-associated traits computed using the pre-intervention phase but identified no significant loci. Finally, we also tested for genotype × diet interaction effects on each of these 25 lifespan-associated traits but identified no significant interaction effects.

**Figure 5:**
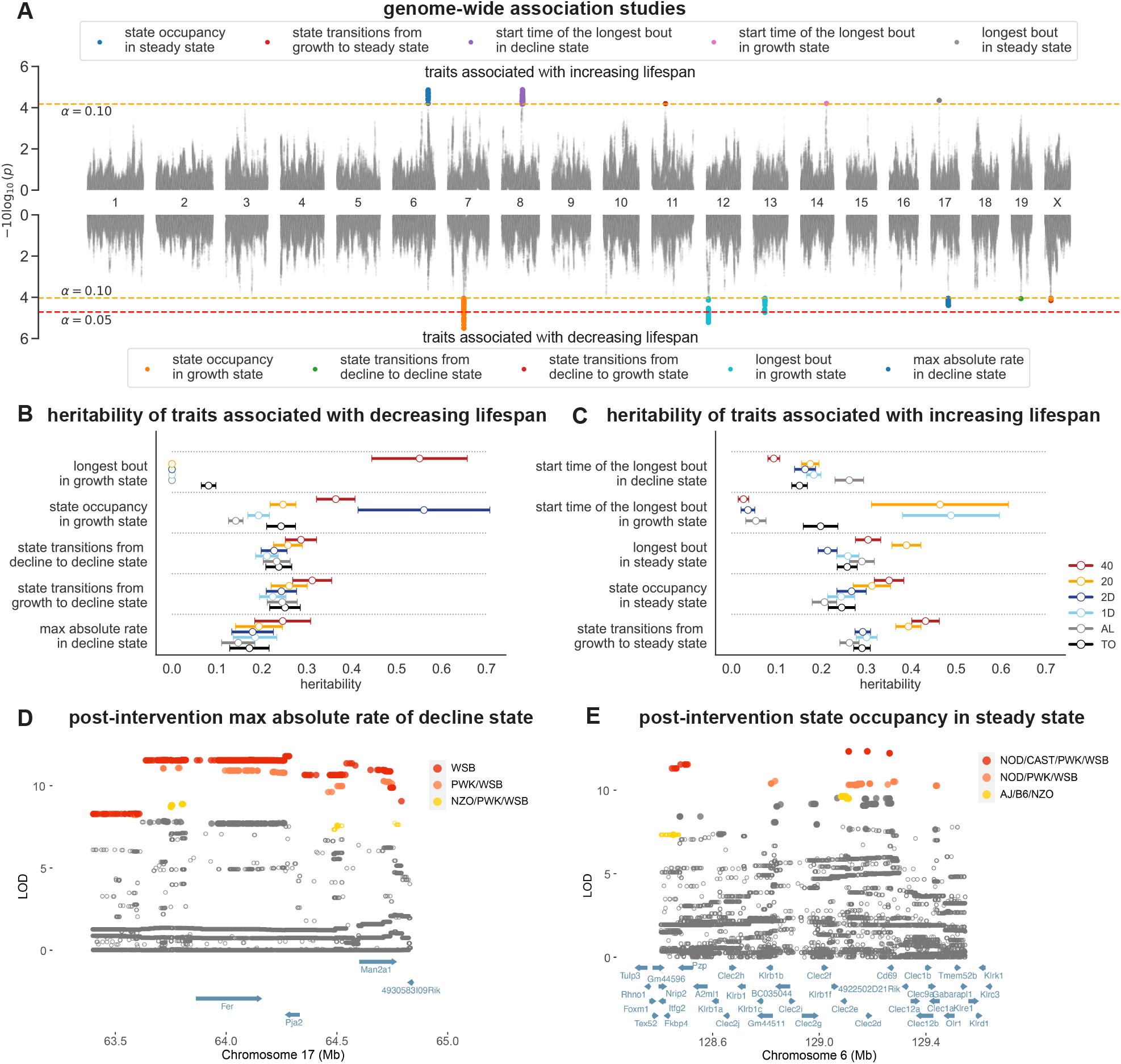
Influence of genetics and environment. (A) Combined Manhattan plot of traits in the post-intervention phase which were either associated with increasing or decreasing lifespan. The red and orange dashed lines represent the group-specific threshold at a false discovery rate of *α*. (B) and (C) Forest plot of the diet-specific heritability and corresponding standard error of different body-weight derived traits, divided into two groups: associated with decreasing lifespan (left) and increasing lifespan (right). Total heritability of the trait is indicated in black. (D) and (E) Fine-mapping loci associated with maximum absolute rate of decline state and steady state homeostasis traits on chromosome 17 and 6, respectively. Solid circles indicate significant variants. Colors denote variants with shared founder allele patterns (FAP). Ranks 1, 2, and 3 by logarithm of the odds (LOD) score are colored red, orange, and yellow, respectively.

We evaluated whether the genetic contribution of the ten lifespan-associated traits differed between various dietary contexts. We quantified the total and diet-dependent heritability for each trait using the GxEMM model, controlling for diet and generation effects. The lifespan-associated traits were less heritable than average body weight post-intervention. Additionally, we observed heritabilities of these traits differed between dietary contexts, with traits typically being more heritable under the 40% CR diet. The higher heritability of these traits in the 40% CR diet can be largely attributed to increased genetic variation relative to other diet groups, suggesting that the dynamics of body weight are tightly regulated under severe calorie restriction. We computed the partial correlation, controlling for diet and cohort effects, between pairs of traits and their genetic correlation using a matrix-variate linear mixed model (Furlotte and Eskin (2015)). We observed the genetic correlation between these traits to be structured, suggesting that the lifespan-associated heritable traits are regulated by shared cellular and molecular processes (**Supplementary Figure 6A**).

To interpret these genetic associations, we fine-mapped each of the 12 significant trait-locus pairs using allele-dosage at both genotyped and imputed variants within a 1-megabase (Mb) window centered at each locus. Our fine-mapping analysis confirmed 7 candidate trait-loci pairs, of which 4 were associated with decreasing lifespan (**Table 1**). We highlight the fine-mapping results of two traits: (a) maximum absolute rate in decline state, which was associated with decreasing lifespan, and (b) steady state homeostasis, which was associated with increasing lifespan (**Figures 5D-5E**). At the chromosome 17 locus associated with *maximum absolute rate of decline*, we grouped fine-mapped variants based on their founder-allele-pattern (FAP) and ranked groups based on the largest LOD score among its constituent variants, to identify variants and haplotypes most likely responsible for the association at this locus (**Methods**). The top FAP group contained variants with minor alleles specific to the WSB strain (**Figure 5B, Supplementary Figure 6B**). All significant variants were located in a genomic interval containing three genes: *Fer, Pja2*, and *Man2a1* (**Figure 5D**). Intersecting the WSB-specific significant variants with regulatory elements assayed across a number of mouse tissues (Cusanovich et al. (2018); Gorkin et al. (2020)), we observed that the strongest variants (*p*-value < 10^−5^) were located within regulatory elements around *Fer* and *Man2a1* genes, active across adipose, intestine, and muscle tissues (**Supplementary Figure 6C**). These observations suggest that this locus likely influenced the rate of loss of body weight by regulating gene expression in tissues directly related to metabolism. Similarly, at the chromosome 6 locus associated with *steady state homeostasis*, the top FAP group contained variants with minor alleles shared by the NOD, CAST, PWK, and WSB strains (**Figure 5E, Supplementary Figure 6D**). All significant variants were located in a genomic interval containing the following genes: *Pzp, A2ml1*, and *Nkrp1-Clr* cluster (**Figure 5E**). The strongest candidate variants were located within regulatory elements at the *Pzp* and *Clec2d* genes, active across most tissues (**Supplementary Figure 6E**).

**Table 1:**
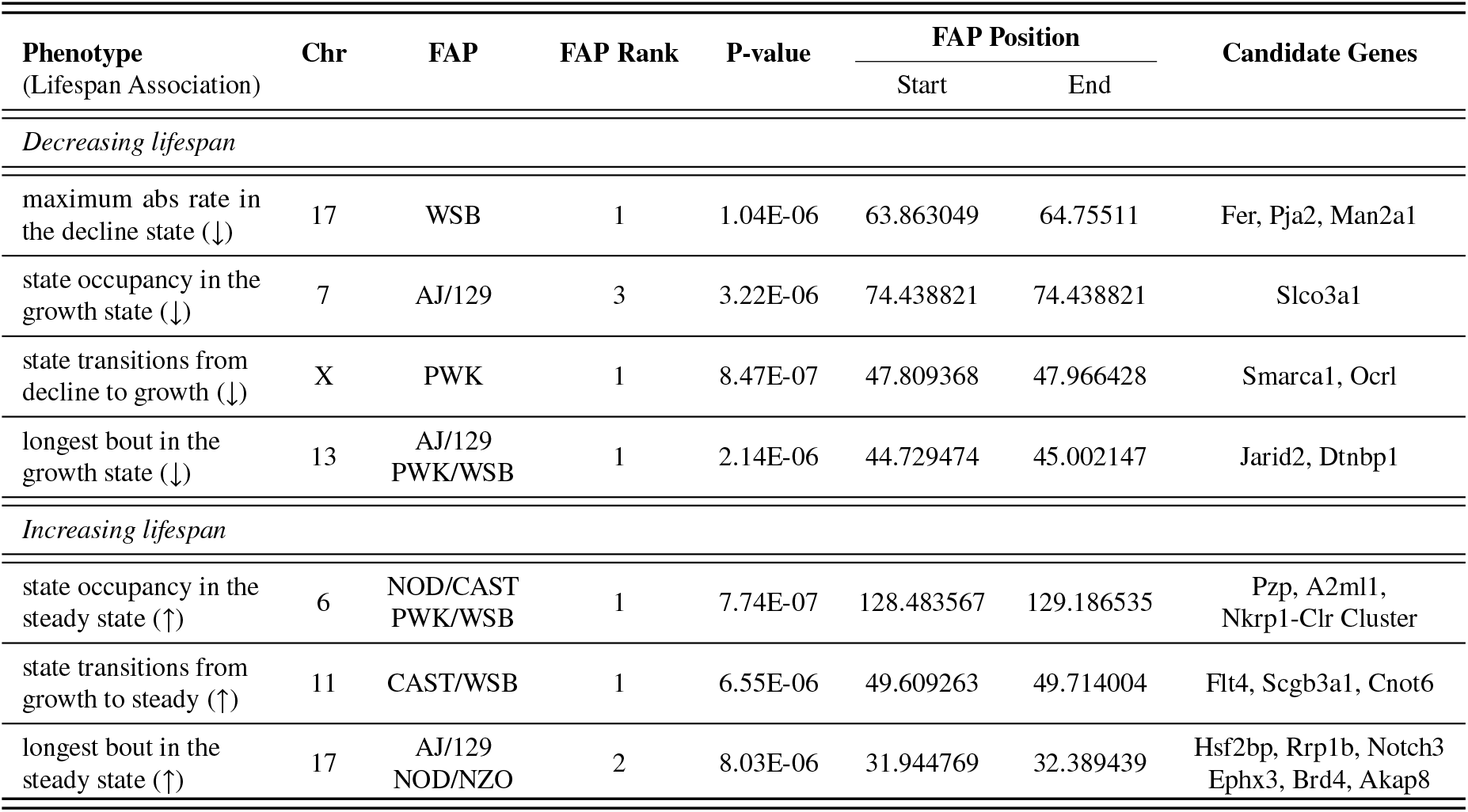
Fine mapping results of body weight-derived phenotypes associated with either increasing or decreasing lifespan in the post-intervention phase.

## Discussion

In this study, we derived novel phenotypes describing the dynamics of body weight measurements taken at high temporal resolution throughout life, evaluated their utility for predicting lifespan, and quantified their genetic determinants. To quantify temporal dynamics in body weight, we developed an autoregressive hidden Markov state space model to extract states representing growth, decline, and maintenance of steady body weight (**Figure 2**). Previous studies investigated age-related trends in body weight from longitudinal measurements using simple parametric models. Due to the non-monotonic nature of body weight dynamics, studies have fit an asymmetrical *inverted-U* pattern either by piece-wise linear regression or polynomial regression and dividing the age-axis into early (growth), mid (steady), and late (decline) life based on the estimated slopes (Wagener et al. (2013)). Such approaches are well suited to capturing long-term trends in mean body weight, but do not capture short-term, transient changes in body weight within each phase such as loss or gain and subsequent stabilization of body weight in response to stressful life events. In contrast, our model of body weight trajectories as a sequence of states allowed for the detection of multiple, transient switches between these states that occur throughout life and captured the continuous nature of body weight fluctuations (i.e., did not require fixed breakpoints of the age axis). Using these inferred states, we derived novel traits that quantify homeostasis of body weight dynamics and adaptation to stress (**Figure 3**). We further demonstrate that these traits are modulated by dietary restriction and predictive of lifespan.

Prior research has demonstrated that body weight during early life predicts lifespan (Miller et al. (2002); Roy et al. (2021)). Similarly, it is well-recognized that diet has an effect on body weight (Fontana et al. (2010); Di Francesco et al. (2018); Longo and Panda (2016); de Cabo and Mattson (2019)). However, no previous studies have quantified the effect of diet on body weight trajectories throughout life and how features derived from such trajectories are associated with lifespan. In our work, we showed that both CR and IF increased the proportion of time spent in a state without dramatic weight loss or gain (steady state). This was not solely due to an inability to gain body weight due to caloric constraints or a physiological lower limit on body weight preventing weight loss. We observed that mice under 1D IF were consuming the same number of calories as ad libitum mice and that mice on CR and IF did spend brief periods of time in both growth and decline states but returned more quickly to steady state compared to ad lib mice. We also demonstrated that time spent in steady state was strongly associated with lifespan at multiple ages, even after conditioning on the effects of diet and body weight (**Figure 4**). Deviation from steady state, captured by the time spent in either growth or decline states, were influenced by diet and phenotyping events, and also associated with lifespan.

One interpretation of deviation from steady state is a perturbation to body weight due to extrinsic stress such as a phenotyping event. This is supported by observation that deviations often coincided with weeks where phenotyping was performed. We hypothesized that the rate of return to steady state is a measure of adaptation to stress. We note that this measure of adaptation to stress is different from *resilience*, a phenotype traditionally used in the field of aging. In biology, *resilience* is often defined as the ability to recover following an acute stress, whereas our measure of adaptation to stress does not require recovery of the original body weight, but rather a return to homeostasis at a new and potentially different body weight (Kirkland et al. (2016)). Our definition of adaptation to stress is more aligned with the definition of resilience in the field of systems dynamics, where the resilience of a system is the rate at which the system converges to an equilibrium state (steady state) after a disturbance (Hirsch and Smith (2006)). Thus, when a mouse returns to steady state following a phenotyping event, it is not required that the mouse regains the body weight lost during the phenotyping event.

While most studies used inbred strains to evaluate the metabolic effects of CR and IF, in this study, we used mice from a genetically diverse outbred stock which enables us to estimate genetic effects on body weight dynamics. Out of 25 lifespan-associated traits, genetic mapping identified 10 traits mapped to 12 genomic loci, including time spent in or away from steady state (**Figure 5**). We estimated diet-specific heritability for each of the 10 lifespan-associated traits and found six had significant differences in heritability between diets. We highlight two traits of interest and their respective associated genetic loci. One trait, *maximum rate of decline in body weight*, was negatively associated with lifespan and fine-mapped to the gene *Man2a1*, with the strongest candidate variants located within regulatory elements that were active in adipose, intestine, and muscle tissues. Mannosidase alpha class 2A member 1 (*Man2a1*) encodes a glycosyl hydrolase that has previously been associated with the pathogenesis of inflammatory bowel disease (Suzuki et al. (2018)), and its inhibition ameliorated rapid body weight decline and symptoms of ulcerative colitis in mouse models. Another trait, *proportion of time spent in steady state*, was positively associated with lifespan and fine-mapped to the gene *Pzp*, where the fine-mapped variants colocalized within a regulatory element active across several tissues relevant to metabolism including adipose and liver tissues. Pregnancy zone protein (*Pzp*), a member of the alpha-2 globulin family of proteins, was recently identified as a key hepatokine regulating factor for fasting-refeeding triggered energy homeostasis through inter-organ cross talk between liver and brown adipose tissue (Lin et al. (2021)).

Our study provides valuable insights into the changes of body weight dynamics. We examine their relationship with age and diet as well as lifespan and genetics. There are some limitations to consider. It is known that body composition and adiposity are also strongly influenced by diet and energy expenditure, and are predictive of lifespan. Future studies would ideally also include dense longitudinal measurements of body composition, food intake, and energy expenditure. Given the importance of body weight changes in response to extrinsic events, a study design that samples body weight more densely around such events (e.g., daily or more often) would allow for a more accurate characterization of adaptation to stress. A measure of how well body weight is regulated in response to stressful events could help identify factors that promote resilience. Additionally, combining sparse temporal sampling of molecular and cellular data (epigenetics, proteomics, metabolomics, and lipidomics) with the dynamics of densely sampled body weight could provide new insight into the mechanisms underlying homeostasis and aging.

## Methods

### Mouse housing, feeding, and body weight measurements

The DO mice were generated by breeding eight founder inbred strains (AJ, B6, 129, NOD, NZO, CAST, PWK, and WSB) to produce an outbred heterozygous population with a random assortment of genetic variation. In this study, 960 DO female mice, sampled at generations 22-24 and 26-28, were enrolled into the study after wean age of 3 weeks old, and maintained at the Jackson Laboratory in Maine. No mice in the study were siblings and maximum genetic diversity was achieved. There were two cohorts per generation for a total of 12 cohorts and 80 animals per cohort. Enrollment occurred in successive quarterly waves starting in March 2016 and continuing through November 2017. Mice were housed in pressurized, individually ventilated cages at a density of eight animals per cage with random cage assignments. They were subject to 12 hours of continuous light and dark cycles beginning at 06:00 AM and 06:00 PM, respectively. Animals exit the study upon death. All animal procedures were approved by the Animal Care and Use Committee at The Jackson Laboratory.

For mice under calorie restriction (CR), food was weighed out for an entire cage of eight. Observation of the animals indicated that the distribution of food consumed was roughly equal among all mice in a cage across diet groups. As the number of mice in each CR cage decreased over time, the amount of food given to each cage was adjusted to reflect the number of mice in that cage. Nonetheless, individual feeding behavior was not controlled among the CR mice and it is possible that some of the variability observed in the CR treatment groups was due to varying degrees of individual caloric intake.

The body weight measurements for this analysis were collated in October 2021, at which point 960 mice (97.35%) had measurements at 6 months, 884 (92.08%) at 12 months, 803 (83.64%) at 18 months, 639 (66.56%) at 24 months, 416 (43.33%) at 30 months, 187 (19.47%) at 36 months, and 43 (4.47%) at 42 months. We included all body weight measures for each mouse up to 1632 days of age (>42 months). We excluded four mice from the downstream analysis (1 from AL group, 2 from 1D group, and 1 from 2D group) because these mice died early in the study and had fewer than seven body weight measurements in total, resulting in 956 mice in our analyses. To obtain a uniform sampling interval for the body weight dynamics, we rounded the time (in days) of measurement of the body weight to the nearest ten. When two body weight measurements were available at the nearest ten, we took the average of these measurements and assigned it to the nearest ten. This strategy of creating a uniform sampling of ten day sampling interval resulted in 4.83% missing values at specific time points. Because the number of missing values is a tiny fraction of the entire database, we used a one-dimensional linear interpolation filter to fill the missing values.

### Autoregressive hidden Markov model

Autoregressive models are based on the idea that the current value of the time-series, *y*_*t*_ measured at time *t*, can be explained as a function of *ℓ* past values, where *ℓ* is the lag-order and determines the number of steps into the past needed to forecast the current values. The simplest form of an autoregressive model can be represented as AR(*ℓ*) which takes the form *y*_*t*_ = *ϕ*_0_ + *ϕ*_1_*y*_*t*−1_ + … + *ϕ*_*l*_ *y*_*t*−*ℓ*_ + *ϵ*_*t*_, where (*ϕ*_0_, …, *ϕ*_*ℓ*_) are the autoregressive coefficients and *ϵ*_*t*_ is the additive white Gaussian noise distributed as 𝒩(0, *σ*^2^) (see **Figure 2A**). A hidden Markov model is a state-space model which is characterized by the hidden states and observations generated by the hidden states. The hidden states are assumed to follow a first-order Markov chain and can only be detected through the observed sequence as they emit observations on varying probabilities. A Gaussian hidden Markov model where *s*_*t*_ ∈ {1, …, *K*} represents latent state at time *t* is specified with an initial probability distribution *π*_0_ ∈ ℝ^*K*×1^, a transition matrix *A* ∈ ℝ^*K*×*K*^ with each *a*_*i*, *j*_ representing the probability of moving from state *s*_*t*−1_ = *i* to state *s*_*t*_ = *j* such that the row sum equals one, and a sequence of observation likelihoods, also called as emission probabilities, each expressing the probability of an observation *y*_*t*_ being generated from state *s*_*t*_ = *i* drawn from a Gaussian distribution 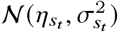 (see **Figure 2A**).

The ARHMM combines an autoregressive model and a hidden Markov model. In an ARHMM, the observations are generated by a few autoregressive time-series models of fixed lag-order, where the switching between these models is controlled by the hidden states which follow a first-order Markov chain. We denote the ARHMM of *l*-th lag-order as ARHMM(*ℓ*) and is defined as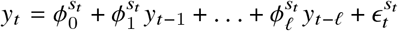, where *s*_*t*_ represents one of the *K* possible latent states, 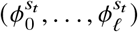 are the autoregressive coefficients corresponding to state *s*_*t*_, and 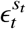 is the state-dependent additive white Gaussian noise distributed as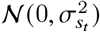. To capture the variation within each latent state, we assumed that the autoregressive coefficients are drawn from a Gaussian distribution of unknown mean and variance given as 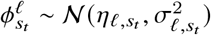, where *ℓ* is the lag-order of the autoregressive model in state *s*_*t*_ (see **Figure 2A**).

The physiological states are represented by the discrete latent states that capture the underlying body weight dynamics at an organismal level. Furthermore, we only considered autoregressive models of lag-order one and set the autoregressive coefficient of the first lag-order to be one, i.e., 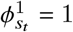, ∀1 ≤ *s*_*t*_ ≤ *K*. Therefore, 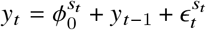, which implies that the body weight at time *t* depends on body weight at time *t* − 1, a state-dependent random variable 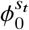 which is drawn from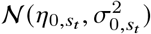, and a state-dependent error which is drawn from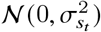. These simplifications made the latent states extracted using the ARHMM interpretable because the physiological states are determined based on the sign and magnitude of the mean of the 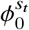 autoregressive coefficient. We implemented the ARHMM using a varaitional inference approach. The details of the derivation are provided in the Appendix.

### Adaptation to stress

We used the posterior probability of steady state as a proxy to measure the rate of recovery because mice spend a significant proportion of their lifespan in the steady state. To compute the rate of recovery, first we identified regions of homeostasis. A region of homeostasis starts when the posterior probability of steady state goes below 0.05, and ends when the probability of steady state stops monotonically increasing or is monotonically increasing and goes beyond 0.95. Thereafter, for each region, we fit a cumulative distribution function of an exponential distribution, which is given as (1 − *e*^−*λx*^), where *λ* is the rate parameter of an exponential distribution. Finally, for any given region of homeostasis longer than thirty days duration, we had at least 3 measurements to estimate *λ* and the value of *λ* that generated the best fit was considered as the rate of recovery.

### Time-varying Cox proportional hazard model

In a traditional Cox model, the hazard ratio is dependent on the covariates and independent of time. In contrast, in an extended Cox proportional model or a time-varying Cox proportional model, the hazard ratio is dependent on time, i.e, the covariates vary with time. The time-varying covariates are classified into two types: internal (dependent on individuals in the study) and external (independent of the individuals in the study). In our proposed model for lifespan analysis, we used body weight as an internal time-varying covariate and the body weight-derived trait measured in a given time-interval bin as an external time-varying covariate. We considered the following time-varying Cox proportional model:

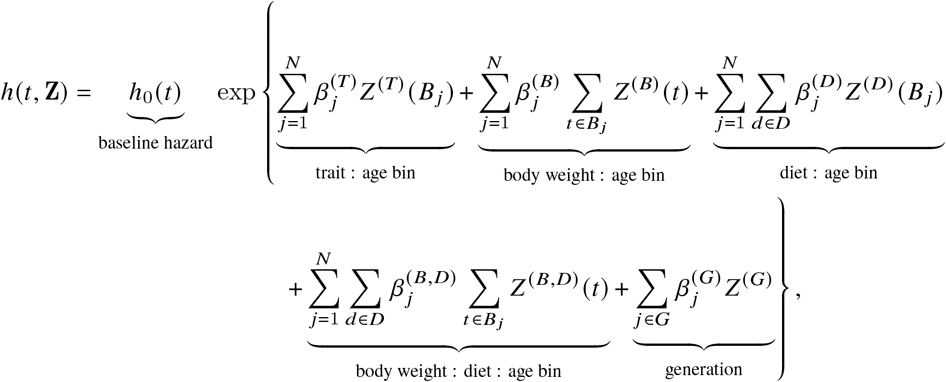

where *h*_0_(*t*) is the baseline hazard, *N* is the number of non-overlapping and continuous age bins represented as {*B*_1_, …, *B*_*N*_}, and 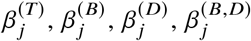, and 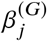 are the effect sizes of (trait : age bin) interaction covariates, (body weight : age bin) interaction covariates, (diet : age bin) interaction covariates, (body weight : diet : age bin) interaction covariates, and generation covariates, respectively. We used the lifelines package implemented in Python to estimate the effect sizes of the body weight-derived trait at every six month interval.

### Genotype measurements

We collected tail clippings and extracted DNA from 954 animals. Samples were genotyped using the 143,259-probe GigaMUGA array from the Illumina Infinium II platform by NeoGen Corp. We evaluated genotype quality using the R package: qtl2. We processed all raw genotype data with a corrected physical map of the GigaMUGA array probes. After filtering genetic markers for uniquely mapped probes, genotype quality, and a 20% genotype missingness threshold, our dataset contained 110,807 markers. Next, examined the genotype quality of individual animals. We found seven pairs of animals with identical genotypes, which suggested that one of each pair was mislabeled. We identified and removed a single mislabeled animal per pair by referencing the genetic data against coat color. Next, we removed a single sample with missingness in excess of 20%. The final quality assurance analysis found that all samples exhibited high consistency between tightly linked markers: log odds ratio error scores were less than 2.0 for all samples. The final set of genetic data consisted of 946 mice. For each mouse, starting with its genotypes at the 110,807 markers and the genotypes of the 8 founder strains at the same markers, we inferred the founders-of-origin for each of the alleles at each marker using the R package: qtl2. This allowed us to test directly for association between founder-of-origin and phenotype (rather than allele dosage and phenotype, as is commonly done in QTL mapping) at all genotyped markers. Using the founder-of-origin of consecutive typed markers and the genotypes of untyped variants in the founder strains, we then imputed the genotypes of all untyped variants (34.5 million) in all 946 mice. Targeted association testing at imputed variants allowed us to fine-map QTLs to a resolution of 1–10 genes.

### Genetic mapping with genotyped markers

For each phenotype, we tested for additive effects between the founder-of-origin at each genotyped variant and the phenotype using the GxEMM model (Wright et al. (2022)), controlling for diet and cohort fixed effects and allowing for genotype-x-diet random effects. Expecting that the set of phenotypes in our genetic analyses were not all independent, we grouped the phenotypes into four groups to determine a group-specific genomewide-level significance threshold. First, we applied a SNP-level false discovery rate of *α* to all the phenotypes within each group. Thereafter, we generated a group-specific *p*-value array consisting of the maximum group-specific *p*-value if the SNP-level null hypothesis is rejected and a randomly drawn group-specific *p*-value if the SNP-level null hypothesis is not rejected. Finally, we applied a false discovery rate of *α* to the *p*-value array of each group separately to obtain a group-specific genome-level significance threshold. This procedure resulted in a conservative and moderate threshold of 1.94 × 10^−5^(at *α* = 0.05) and 8.99 × 10^−5^(at *α* = 0.10), respectively, for those phenotypes associated with decreasing lifespan in the post-intervention phase. Similarly, we obtained a moderate threshold of 6.75 × 10^−5^(at *α* = 0.10) for those phenotypes associated with increasing lifespan in the post-intervention phase. Pre-intervention phenotypes, either associated with increasing or decreasing lifespan, did not result in any significant variants at FDR of *α* = 0.05 and *α* = 0.1.

### Genetic fine-mapping with founder-allele-patterns

To more precisely fine-map the genomic interval of each QTL, we imputed all SNPs and insertion-deletion variants from the fully sequenced DO founders for a 5 Mb interval centered at the lead genotype-marker. For each imputed variant, we identified the founder-of-origin for the major and minor allele. To illustrate the process, consider the bi-allelic A/G variant. If allele A was specific to founders AJ, NZO, and PWK, and allele G was specific to the other five founders, then we assigned A to be the minor allele and defined a founder-of-origin allele pattern (FAP) of AJ/NZO/PWK for this variant. We identified variants and founder haplotypes most likely responsible for the association at each locus by grouping variants based on their FAP and ranked groups based on the largest logarithm of odds (LOD) score among its constituent variants. We hypothesized that the functional variant(s) responsible for trait-specific variation were among those in the lead FAP group because they exhibit the strongest statistical association and it is unlikely any additional variants are segregating in this genomic interval beyond those identified in the full genome sequences of the eight founder strains. By focusing on the FAP groups with the largest LOD scores, we significantly reduced the number of putative causal variants, while representing the age- and diet-dependent effects of these loci in terms of the effects of its top FAP groups. We further narrowed the number of candidates by intersecting the variants in top FAP groups with functional annotations (e.g., gene annotations, regulatory elements, tissue-specific regulatory activity, etc.). This procedure identified candidate regions containing 1 − 3 genes.

## Code Availability

The code for extracting novel body weight-derived traits using an autoregressive hidden Markov model, as well as the subsequent lifespan and genetic analyses of these traits, is publicly available through the following GitHub repository: https://github.com/calico/do_bwd. The repository also contains trained models, processed datasets, and scripts that can be utilized to train models from scratch and generate processed datasets.

## Acknowledgements

We thank the JAX Nathan Shock Center Animal and Phenotyping core team for their assistance with animal husbandry, data collection, and curation. We would also like to thank Eugene Melamud for helpful feedback and discussion. This work was funded by Calico Life Sciences LLC.

## Supplementary Tables

**Table 1:** Estimated parameters of the variational inference-based autoregressive hidden Markov model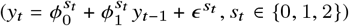.

| State | Initial Probability | Transition Probability from State X |  |  | AR(0) | Error |
| --- | --- | --- | --- | --- | --- | --- |
|  |  | P(DS\|X) | P(SS\|X) | P(GS\|X) |  |  |
| Decline (X = DS) | $\pi_0 = 0.42$ | 0.42 | 0.21 | 0.37 | $\phi_0^0 \sim \mathcal{N}(-1.6, 9.1 \times 10^{-4})$ | $\epsilon^0 \sim \mathcal{N}(0, 2.00)$ |
| Steady (X = SS) | $\pi_1 = 0.07$ | 0.07 | 0.93 | $\approx 0.00$ | $\phi_0^1 \sim \mathcal{N}(+0.1, 4.4 \times 10^{-6})$ | $\epsilon^1 \sim \mathcal{N}(0, 0.64)$ |
| Growth (X = GS) | $\pi_2 = 0.14$ | 0.14 | 0.12 | 0.74 | $\phi_0^2 \sim \mathcal{N}(+1.4, 3.4 \times 10^{-5})$ | $\epsilon^2 \sim \mathcal{N}(0, 1.83)$ |

**Table 2:**
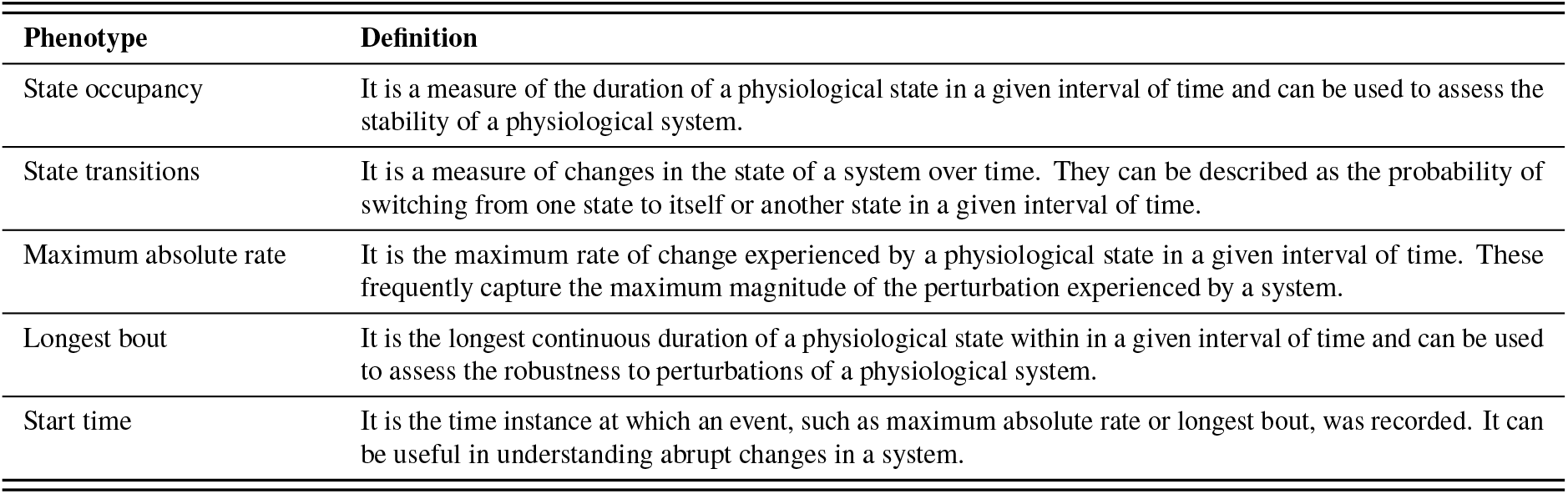
Body weight-derived traits and definitions.

| Phenotype | Definition |
| --- | --- |
| State occupancy | It is a measure of the duration of a physiological state in a given interval of time and can be used to assess the stability of a physiological system. |
| State transitions | It is a measure of changes in the state of a system over time. They can be described as the probability of switching from one state to itself or another state in a given interval of time. |
| Maximum absolute rate | It is the maximum rate of change experienced by a physiological state in a given interval of time. These frequently capture the maximum magnitude of the perturbation experienced by a system. |
| Longest bout | It is the longest continuous duration of a physiological state within in a given interval of time and can be used to assess the robustness to perturbations of a physiological system. |
| Start time | It is the time instance at which an event, such as maximum absolute rate or longest bout, was recorded. It can be useful in understanding abrupt changes in a system. |

## Supplementary Figures

**Supplemental Figure 1:**
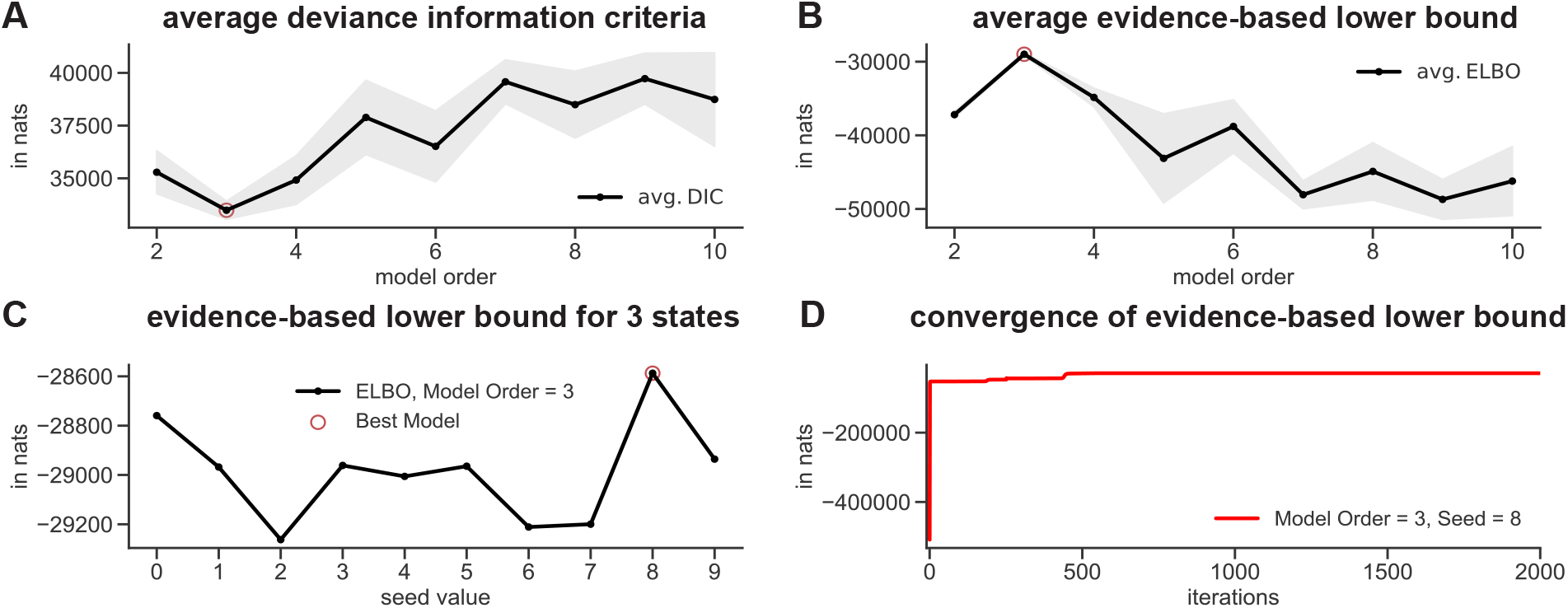
Variation inference-based autoregressive hidden Markov model (VI-ARHMM). (A) The mean value of the deviance information criteria (DIC) and standard error computed on the validation set for different model orders *K* ∈ {2, …, 10} at ten different random state initializations. Model order *K* = 3 resulted in the smallest DIC. (B) The mean value of the evidence-based lower bound (ELBO) and standard error computed on the training set for different model orders at ten different random state initializations. Once again, model order *K* = 3 resulted in the highest ELBO. (C) The ELBO of the training data for model order *K* = 3 at different random state seed values. The highest ELBO was obtained at seed value of 8 (best model). (D) The VIARHMM approach showed convergence of the ELBO for the best model.

**Supplemental Figure 2:**
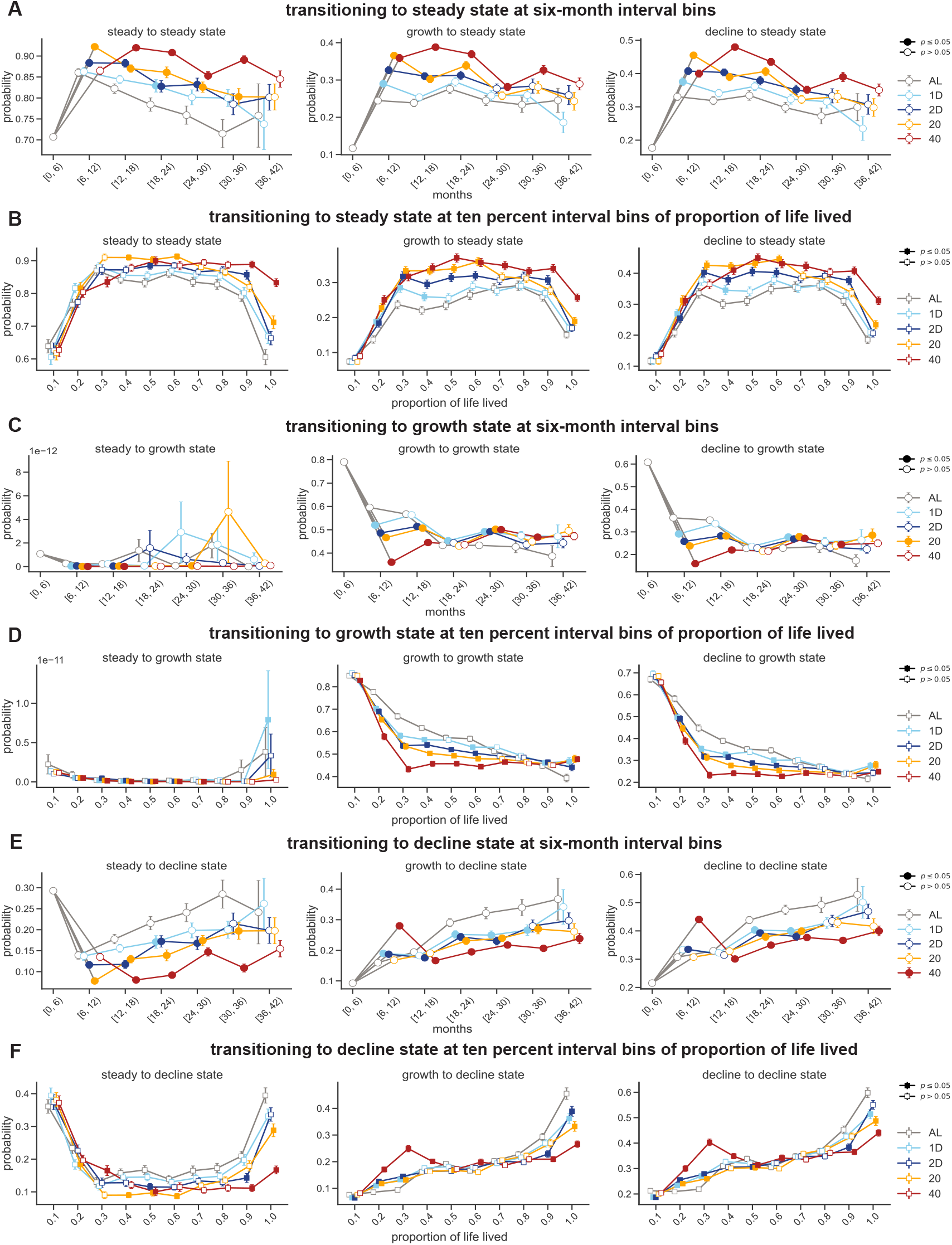
Influence of diet and age on the state stability. (A) and (B) Transitioning into steady state at six month and ten percent of proportion of life lived interval bins, respectively. (C) and (D) Transitioning into growth state at six month and ten percent of proportion of life lived interval bins, respectively. (E) and (F) Transitioning into decline state at six month and ten percent of proportion of life lived interval bins, respectively. In (A)-(F), solid squares or circles indicate *p*-values < 0.05, where the *p*-values were obtained by performing a Mann-Whitney test between a diet group and the AL group conditioned at the same interval bin.

**Supplemental Figure 3:**
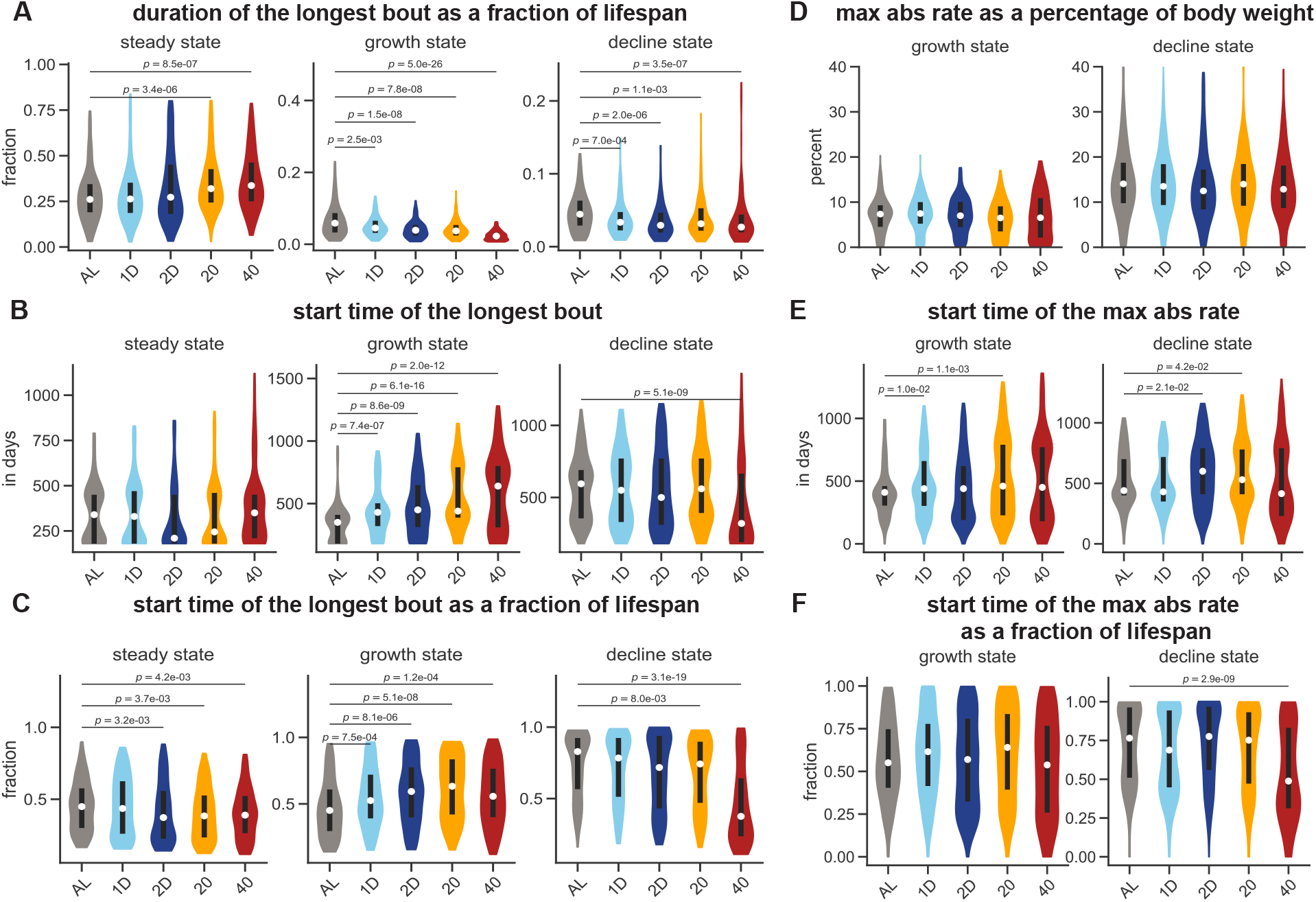
Robustness and perturbations due to body weight homeostasis. (A) Fraction of life spent in the longest continuous bouts of growth, steady, and decline states. (B) The start time of the longest bout in growth, steady, and decline states. (C) Normalized time at which the start time of the longest bout were recorded. (D) Percentage of body weight gained and lost at the time of the maximum absolute rates of growth and decline states, respectively. (E) The time (in days) at which the maximum absolute rates of growth and decline states were recorded. (F) Normalized time at which the maximum absolute rates of growth and decline states were recorded.

**Supplemental Figure 4:**
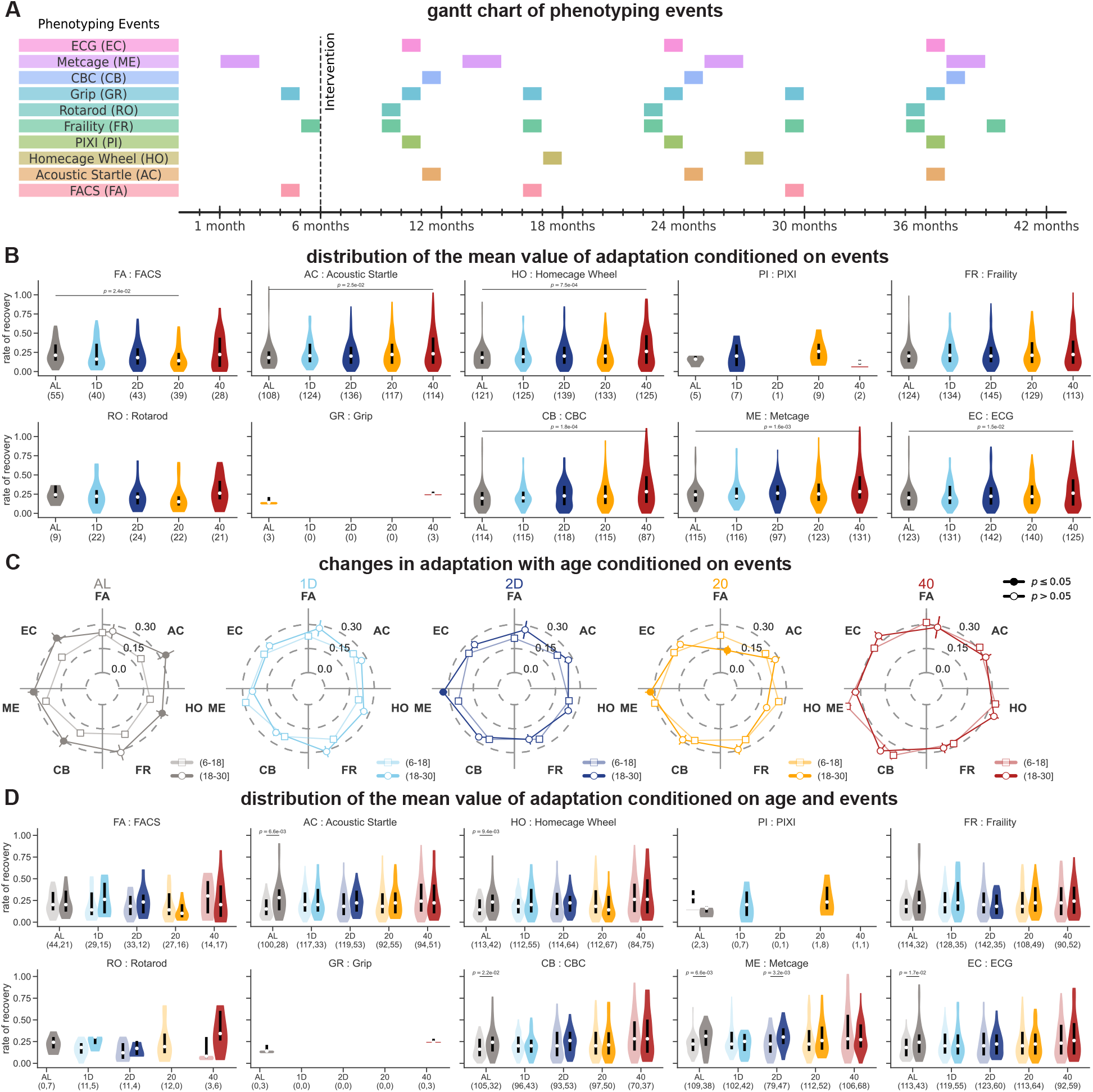
Adaptation to stress is influenced by diet and age. (A) A gantt chart of the different phenotypic assays a mouse undergoes in its lifespan. The approximate duration it takes to assay all mice for a particular assay is indicated using a color bar corresponding to the phenotypic assay. (B) Violin plots of the average values of the rate of recovery, conditioned on the phenotypic assay. (C) Radar charts of the average value and the standard-errors of the rate of recovery conditioned on phenotypic assay for each diet at two non-overlapping age-bins.(D) Violin plots of the average values of the rate of recovery, conditioned on the phenotypic assay and time interval. Intervals [6–18) and [18–30) months are represented in lighter and darker shades, respectively. In (B) and (D), the number of mice for which a perturbation event was registered following a phenotypic assay in the post-intervention phase is mentioned in the parenthesis below the diet. Only a small number of perturbation events were registered after PIXI, rotarod, and grip strength phenotypic assays.

**Supplemental Figure 5:**
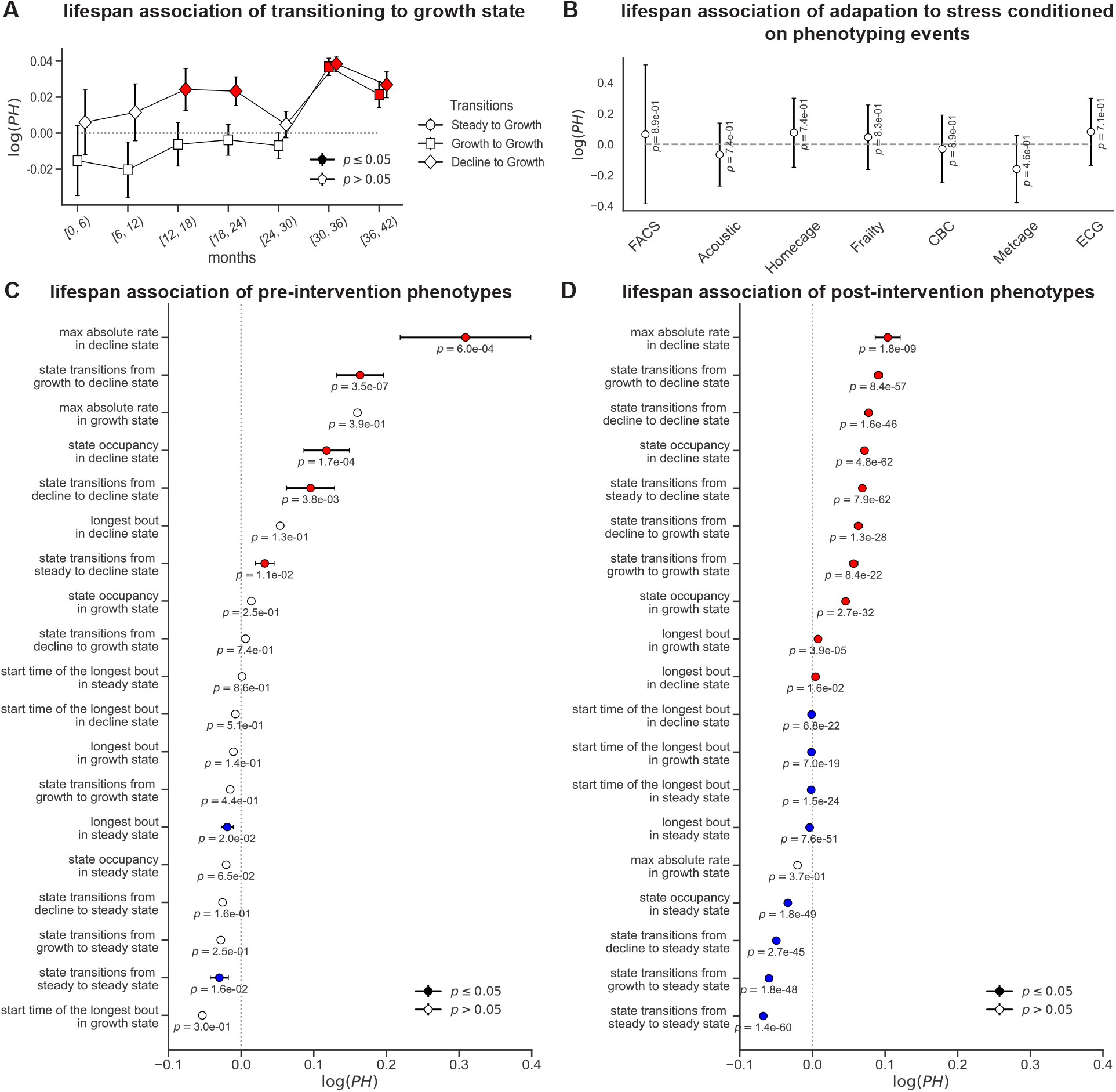
Body weight-derived traits and its association with lifespan. (A) Effect size and standard error of transitioning to growth state and its association with lifespan at every six-month interval bins. (B) Effect size and standard error of the mean value of adaptation to stress conditioned on the phenotyping event and its association with lifespan. (C) and (D) Effect sizes and standard errors of the pre- and post-intervention body weight-derived traits that are associated with either increasing or decreasing lifespan. The traits are arranged in descending order of the effect sizes. In (A)-(D), lifespan associations which were statistically significant (*p*-value < 0.05) were indicated with solid colors. If the effect size of the trait lies above or to the right of the dashed gray line (red filled markers), then an increase in the value of the trait decreases lifespan. Conversely, if it lies below or to the left the dashed gray line (blue filled markers), then an increase in the value of the trait increases lifespan.

**Supplemental Figure 6:**
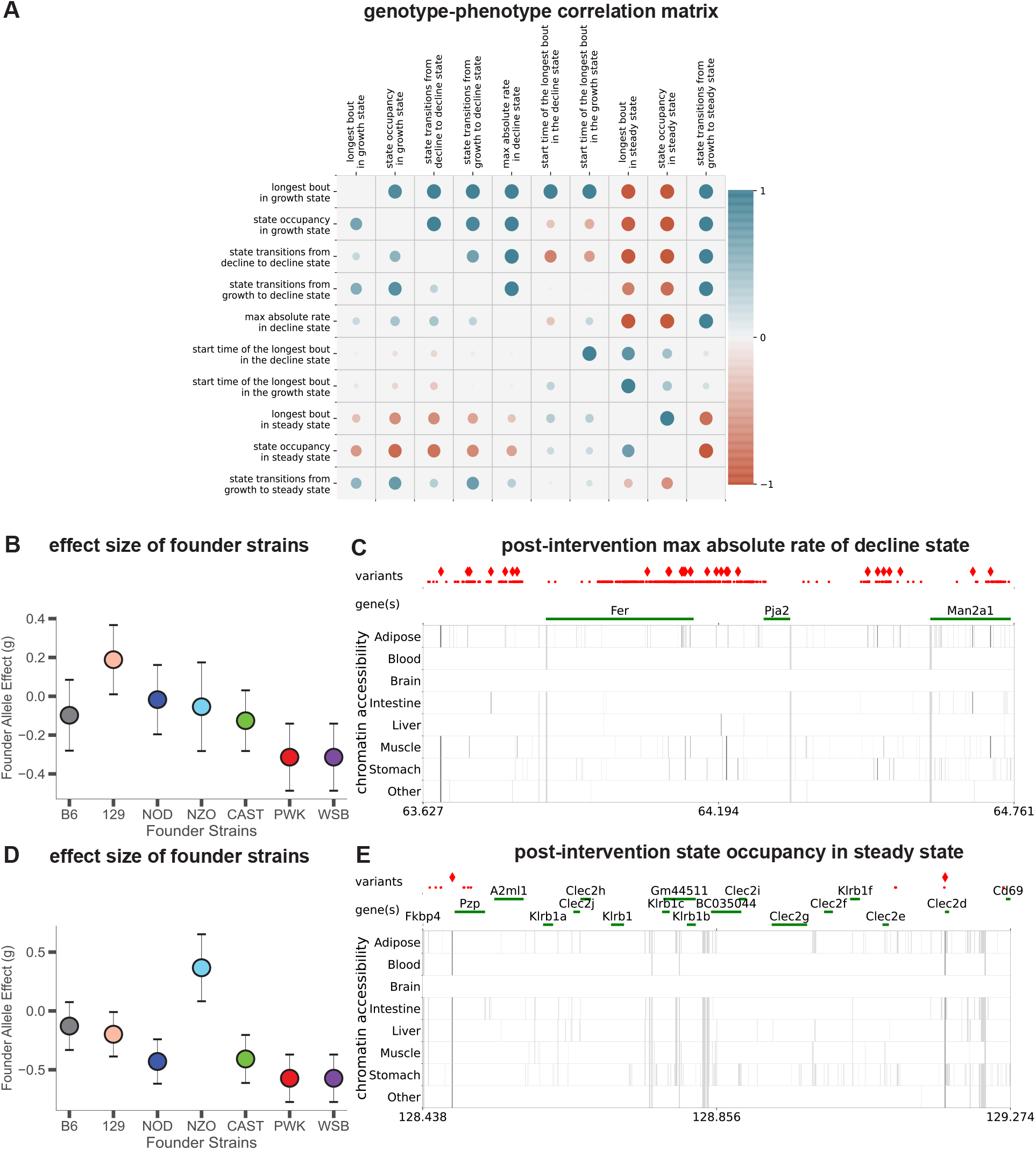
Phenotypic-genotypic correlations and evidence supporting QTL. (A) Pairwise genetic (upper triangle) and phenotypic (lower triangle) correlations of the body weight-derived traits in the post-intervention phase. (B) and (D) Effect size of founder allele strains observed at the variant with the highest LOD score for max absolute rate of decline and steady state occupancy, respectively, in the post-intervention phase. (C) and (E) Significant variants, colored by their founder allele pattern (FAP) group and the tissue-specific activity of regulatory elements near these variants (shown in gray). Significant variants that lie within regulatory elements are highlighted as diamonds, and regulatory elements that contain a significant variant are highlighted in black.

## Appendix

### Variational Inference-based Autoregressive Hidden Markov Model

960 diversity outbred female mice were subject to 5 dietary interventions. The body weight of each mouse was measured each week (approximately) starting from one month of age. Using these high resolution temporal measurements, we wish to learn interesting physiological and developmental stages in the life of a mouse. To identify the different physiological and development stages in the life of a mouse, we develop a variational inference-based auto-regressive model.

- If *M* denotes the number of mice enrolled in the study, we represent the body weight trace of the *m*-th mouse as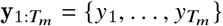, where *T*_*m*_ is the total number of body weight measurements for the *m*-th mouse and the body weight measurements are sorted based on the age of the mouse at the time of the measurement.
- Let 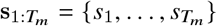 denote a sequence of hidden physiological states corresponding to the body weight measurements 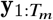, generated by a first-order Markov process, where *s*_*i*_ ∈ {1, …, *K*} and *K* is the total number of physiological states.
- Let ***π*** denote the initial probability vector of the first-order Markov process where,

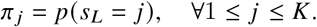
- Let **A** denote the state transition matrix where transition between states is governed by Markov chain whose realizations take on values {1, …, *K*} and the elements of the state transition matrix are given as:

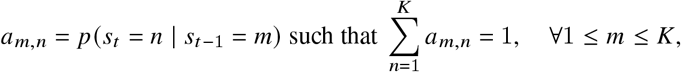

where *a*_*m*,*n*_ is the probability of transitioning from *s*_*t*−1_ = *m* to *s*_*t*_ = *n*.
- The dynamics of the body weight measurements at a given physiological state *s*_*t*_ can be defined using an auto-regressive model as:

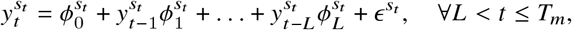

where *L* is the lag order, 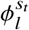 is the auto-regressive coefficient for the *l*-th lag order, and 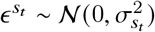 is zero mean additive white Gaussian noise.
- We assume that the mouse-specific parameters are unobserved and propose a hierarchical model on the dynamics parameters. If ***ϕ***_*l*_ = [*ϕ*_*l*,1_, …, *ϕ*_*l*,*K*_]^T^ denotes a vector of autoregressive coefficients for the *l*-th lag order, then ***ϕ***_*l*_ ∼ 𝒩(***η***_*l*_, **Σ**_*l*_), where ***η***_*l*_ = [*η*_*l*,1_, …, *η*_*l*,*K*_]^T^ and **Σ**_*l*_ is a diagonal covariance matrix with diagonal elements 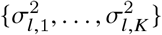. For notation convenience, we concatenate all auto-regressive coefficients into a long vector 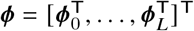.

The joint likelihood can be written as:

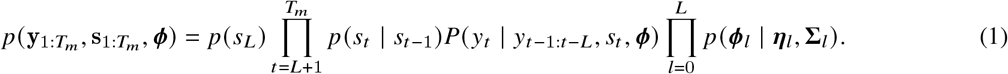

The latent variables 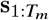 and ***ϕ*** are not observed. Therefore, we integrate the latent variables and rewrite the complete log-likelihood as:

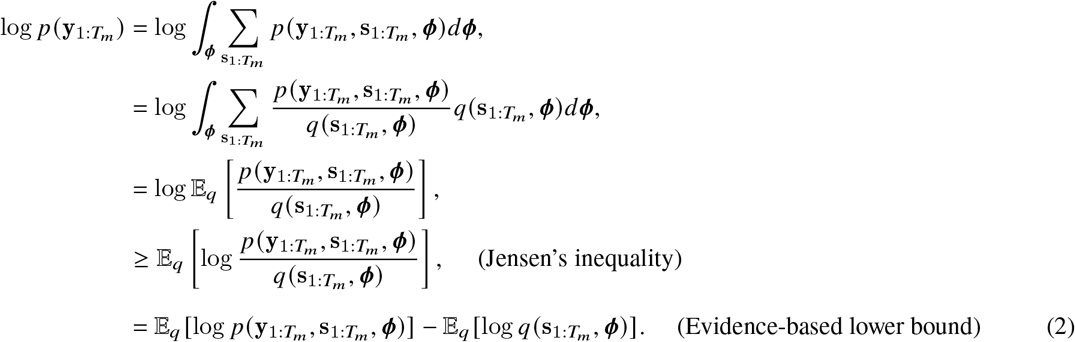

The log-likelihood is maximized when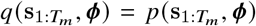. However, the joint probability of 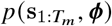 is intractable. The goal of variational inference algorithm is to find the distribution 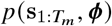 from the chosen variational family that maximizes the lower bound to the log marginal likelihood. We want to optimize the evidence-based lower bound (ELBO) over a chosen space of variational distributions, to find the variational distribution closest to the true posterior distribution 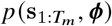 . Therefore, we replace 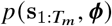 with 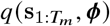 such that 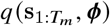 is tractable and 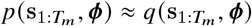. Further, to perform this optimization, we assume that (a) the states and the autoregressive coefficients are independent, (b) autoregressive coefficients are independent, and (c) the transition between states is governed by a first-order Markov process. Under these assumptions,

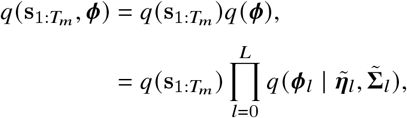

where 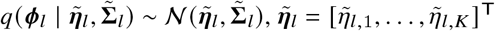 is the mean, 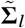 is a diagonal covariance matrix with diagonal elements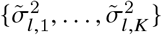,

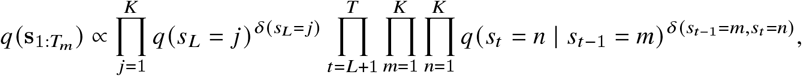

and *δ*(·) acts as a masking function which is set to one if the condition is true else zero. We expand the terms in the ELBO expression in (2) and rewrite it as:

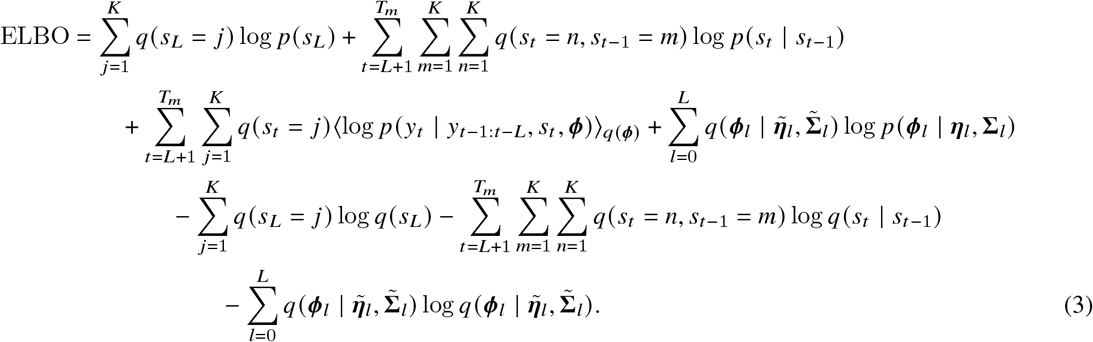

The above expression can be further simplified as:

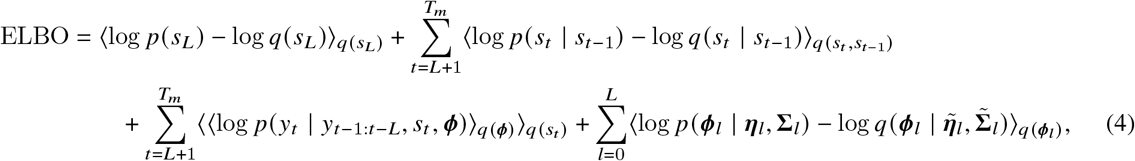

where ⟨·⟩_*q*(·)_ is the expectation with respect to *q* (·). Furthermore, based on the normality assumptions made on 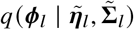, we get

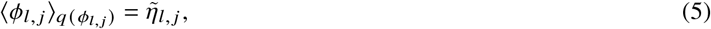

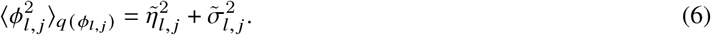

Substituting (5) and (6) in (4) gives the complete expression of ELBO which is given as:

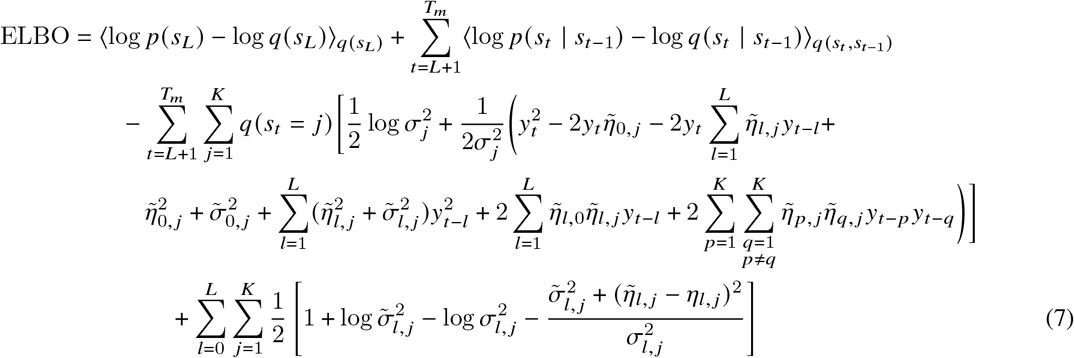

Maximizing the variational parameters of the ELBO by keeping the model parameters fixed gives us the variational E-step and M-step, and maximizing the model parameters of the ELBO by keeping the variational parameters fixed gives us the M-step. These steps are repeated until the convergence criteria is met.

### Variational Bayes E-Step

In the variational Bayes E-step, the mouse-specific and model parameters are fixed and the ELBO is maximized over the mouse-specific variational marginal and joint probabilities. We used the forward-backward algorithm to compute the marginal and joint variational probabilities *q* (*s*_*t*_ = *j*) and *q* (*s*_*t*_ = *n, s*_*t*−1_ = *m*), respectively. The compact expression of ELBO in (2) can be rewritten in a functional form as:

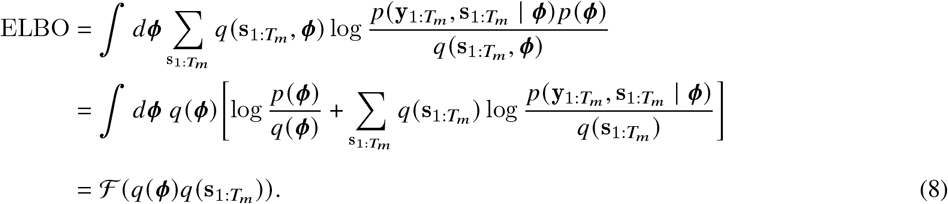

Taking the functional derivative of (8) with respect to 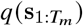 gives:

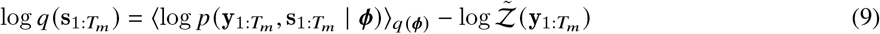

where 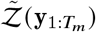 is a normalization constant. The normalization constant enables the feasibility of computing *q* (*s*_*t*_ = *j*) and *q* (*s*_*t*_ = *n, s*_*t*−1_ = *m*) using forward-backward algorithm. Unlike the hidden Markov model which requires the joint log-likelihood, the re-parameterized implementation of the forward-backward algorithm requires the expectation of the partial log-likelihood. Moreover, based on the assumptions made on the variational parameters, we can easily compute the closed-form expression of the expectation of the partial log-likelihood with respect to *q* (***ϕ***) as follows:

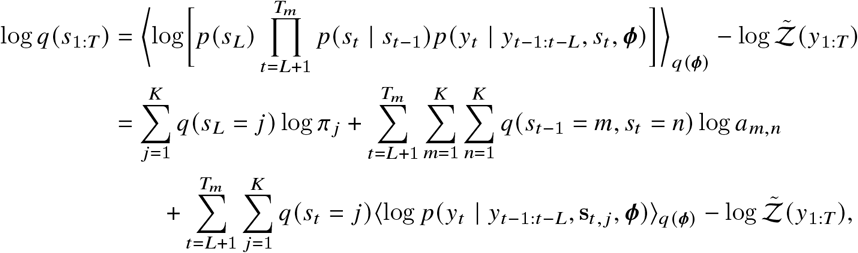

where

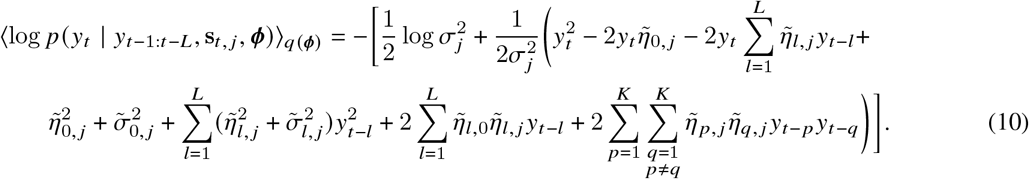

The output of the forward-backward algorithm generates forward and backward probabilities, which are used to compute the variational marginal and joint probabilities for a specific mouse. The main advantage of the reparameterization step is that we can avoid computing the partial derivatives of the marginal and joint probabilities, which are often computationally intensive to solve.

### Variational Bayes M-Step

In the variational Bayes M-step, the mouse-specific variational proabilities and model parameters are fixed and the ELBO is maximized over the variational model parameters.

- Variational initial probability *q* (*s*_*L*_): To get *q* (*s*_*L*_), we solve the following optimization problem:

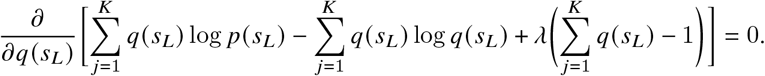 On solving, we get:

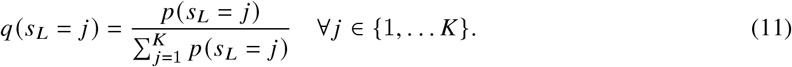
- Variational transition probability *q* (*s*_*t*_ | *s*_*t*−1_): To get *q* (*s*_*t*_ | *s*_*t*−1_), we solve the following optimization problem:

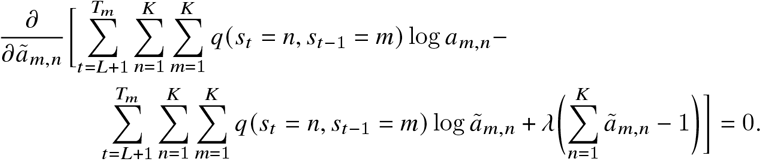 On solving, we get:

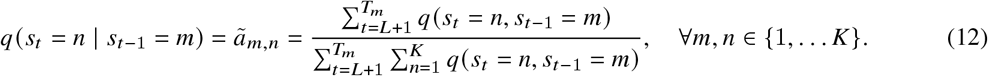
- Variational variance 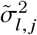:

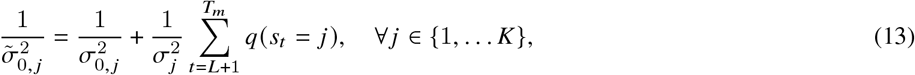

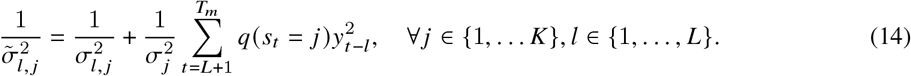
- Variational mean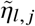:

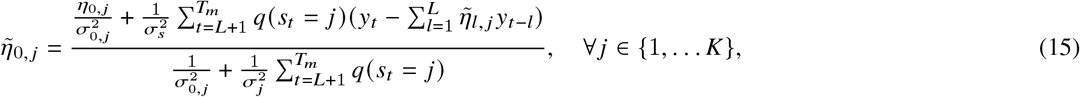

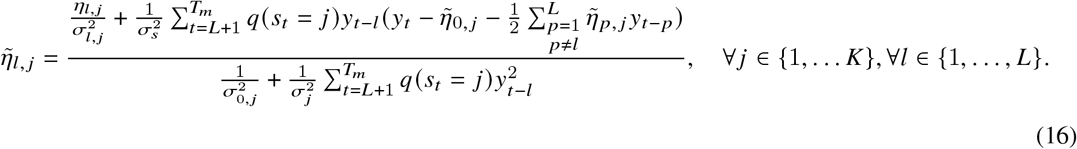

### M-Step

In the M-step, the mouse-specific variational parameters are fixed and the ELBO is maximized over model parameters. We used subscript *i* to indicate the variational parameters from the *i*-th mouse.

- Initial probability *p* (*s*_*L*_):

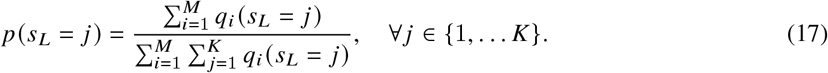
- Transition probability *p* (*s*_*t*_ | *s*_*t*−1_):

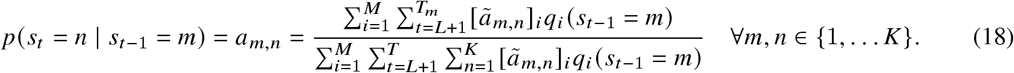
- Error variance 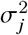

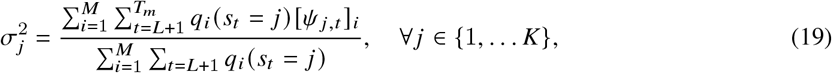

where

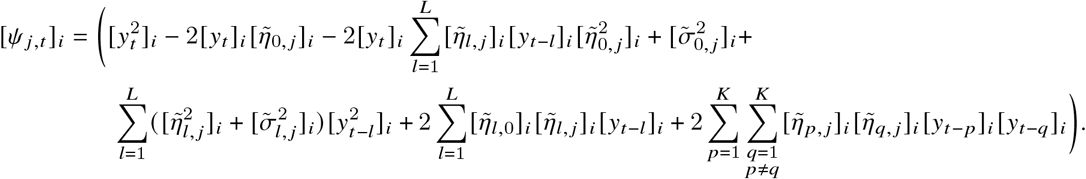
- Auto-regressive coefficient parameters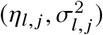:

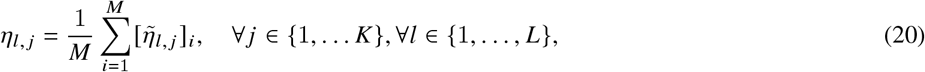

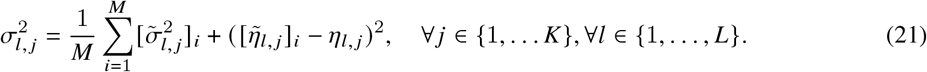

### Model Selection

We used deviance information criteria (DIC) to select the model order for the variational-inference based auto-regressive hidden Markov model (Spiegelhalter et al. (2002)). The DIC combines model complexity and fit, where the model complexity is obtained by taking difference between the posterior mean of the deviance and the deviance calculated at the posterior mean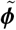, and fit is obtained from the log-likelihood computed at the posterior mean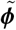. The DIC is expressed as:

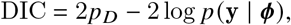

where

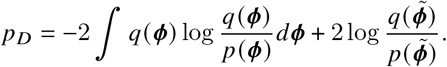

Based on the assumptions made about the model and variational parameters,

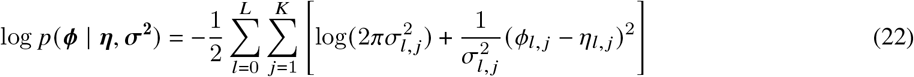

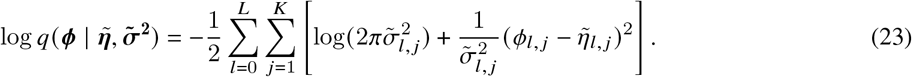

Using (22) and (23), *p*_*D*_ can be simplified as:

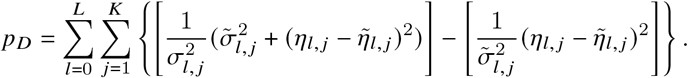

We computed the deviance information criteria at ten different seed values for each model order. The model order with the smallest deviance information criteria is considered as the best model order. To select the best seed value at a given model order, we computed the ELBO and choose the seed value with the highest ELBO.

## Notes

### Competing Interest Statement

GVP, ZC, KW, ADF, VJ, and AR were employees of Calico Life Sciences, LLC during the time of their contribution to the study. The authors declare no other competing interests.

